# Linalool acts as a fast and reversible anesthetic in *Hydra*

**DOI:** 10.1101/584946

**Authors:** Tapan Goel, Rui Wang, Sara Martin, Elizabeth Lanphear, Eva-Maria S. Collins

## Abstract

The ability to make transgenic *Hydra* lines has opened the door for quantitative *in vivo* studies of *Hydra* regeneration and physiology. These studies commonly include excision, grafting and transplantation experiments along with high-resolution imaging of live animals, which can be challenging due to the animal’s response to touch and light stimuli. While various anesthetics have been used in *Hydra* studies over the years, they tend to be toxic over the course of a few hours or their long-term effects on animal health have not been studied. Here we show that the monoterpenoid linalool is a useful anesthetic for *Hydra*. Linalool is easy to use, non-toxic, fast acting, and reversible. It has no detectable long-term effects on cell viability or cell proliferation. We demonstrate that the same animal can be immobilized in linalool multiple times at intervals of several hours for repeated imaging over 2-3 days. This uniquely allows for *in vivo* imaging of dynamic processes such as head regeneration. We further directly compare linalool to currently used anesthetics and show its superior performance. Because linalool, which is frequently utilized in perfumes and cosmetic products, is also non-hazardous to humans, it will be a useful tool for *Hydra* research in both research and teaching contexts.

## Introduction

Abraham Trembley’s careful and systematic studies on *Hydra* regeneration, published in his *Memoires* in 1744, brought this freshwater cnidarian into the spotlight of biological research (Lenhoff & Lenhoff 1986). *Hydra* is an optically transparent polyp a few millimeters in length. It consists of a hollow cylindrical body column with a head on one end, consisting of a ring of tentacles and a dome-shaped hypostome, and an adhesive basal disk on the other end. *Hydra* is composed of only a small number of cell types originating from three (ectodermal, endodermal and interstitial) stem cell lineages (Bode 1996). This anatomical simplicity, continuous cell turnover in the adult (Campbell 1974), and the ability to regenerate from small fragments of the body column or even from aggregates of cells (Gierer et al. 1972; Shimizu et al. 1993) render *Hydra* a powerful system for studies of development (Steele 2002), stem cell biology (David & Murphy 1977; Bosch 2009), and regeneration (Bosch 2007; Galliot et al. 2018; Cochet-Escartin et al. 2017; Petersen et al. 2015). Furthermore, *Hydra* has a relatively simple nervous system (Burnett & Diehl 1964; Bode et al. 1973), consisting of a few thousand cells (David 1973) that are organized in three neuronal networks (Dupre & Yuste 2017), making it an attractive system to study neuron development (Noro et al. 2019; Koizumi 2002) and neuronal control of behavior (Han et al. 2018; Dupre & Yuste 2017).

Exploiting *Hydra*’s patterning processes and regenerative abilities via sophisticated excision and grafting studies has been a mainstay of *Hydra* research since Trembley’s original experiments. This “cut-and-paste” approach has provided fundamental insights into *Hydra* biology. For example, the excision and subsequent threading of body column rings onto fishing line allowed researchers to probe questions about oral-aboral polarization (Ando et al. 1989). Grafting of hypostomes into body columns showed that the tip of the hypostome acts as head organizer (Browne 1909; Yao 1945) long before the head organizer was biochemically analyzed (Bode 2012). Transplantation experiments were used to characterize the properties and dynamics of head inhibition (MacWilliams 1983) and estimate the length scales of head activation and inhibition (Technau et al. 2000), which helped validate the Gierer-Meinhardt model of axial patterning (Gierer & Meinhardt 1972) long before *in vivo* visualization of cells or proteins was possible in *Hydra*.

However, despite its many advantages, *Hydra* has not become a mainstream model organism like fruit flies or nematodes due to the lack of genetic tools so readily available in these organisms. This has changed in the last decade, with access to a fully assembled *Hydra* genome (Chapman et al. 2010), single cell RNAseq data (Siebert et al. 2018), and the development of molecular tools that allow for the generation of transgenic lines (Juliano et al. 2014; Glauber et al. 2015; Wittlieb et al. 2006). Because of these tools, numerous recent studies have been able to address longstanding open questions that could not previously be answered. For example, the recent creation of a transgenic line expressing GCaMP6s in the interstitial lineage allowed visualization of neural activity in real time in freely behaving animals and led to the discovery of multiple discrete networks of neurons linked to specific behaviors (Dupre & Yuste 2017). Transgenic animals have also enabled quantitative biomechanics studies to conclusively settle key biological questions, such as the mechanism driving cell sorting during regeneration from cell aggregates (Cochet-Escartin et al. 2017) and the functioning of the *Hydra* mouth (Carter et al. 2016).

As research in the field continues to dig deeper into such questions in the living animal, studies will require ever more precise and repeatable manipulations of the animal, high resolution live imaging, or a combination of the two to fully exploit transgenic strains and other new technologies. These experimental approaches are challenged by the fact that the animal is in a continuous dynamic state of extension-contraction and responds rapidly to stimuli such as touch and light. Therefore, a reversible way of slowing or preventing the animal’s movements would greatly facilitate a wide range of experiments. The search for a reliable and reversible relaxant in *Hydra* has driven the field to try an array of compounds, with the most prominent ones being urethane (Macklin 1976; Benos et al. 1977; Münder et al. 2013; Takahashi & Hamaue 2010; Buzgariu et al. 2018), heptanol (Smith et al. 2000; Rentzsch et al. 2005), and chloretone (Badhiwala et al. 2018; Lommel et al. 2017; Loomis 1955; Kepner & Hopkins 1938). Urethane and heptanol have broad effects on *Hydra.* Urethane reverses the transepithelial potential, causing adverse effects upon several hours of exposure (Macklin 1976). Heptanol blocks epithelial gap junction communication in the body column (Takaku et al. 2015). Chloretone is reportedly nervous-system specific, but *Hydra* was observed to develop tolerance to the anesthetic within hours of exposure (Kepner & Hopkins 1938). Thus, existing anesthetics have serious limitations and there is an urgent need for an alternative that reliably immobilizes *Hydra* without causing tolerance or adverse health effects.

Here we report on linalool as a novel, safe and fully reversible anesthetic for *Hydra*. Linalool is a monoterpenoid alcohol found in flowers and frequently used in cosmetic products (Aprotosoaie et al. 2014). It has previously been shown to have anesthetic or sedative activity in a range of other model systems, including mice (Linck et al. 2009), catfish (Heldwein et al. 2014), and flatworms (Boothe et al. 2017). Linalool exists in two enantiomeric forms which are known to have different pharmacological effects. In humans, the (S)-enantiomer causes an increase in heart rate while the (R)-enantiomer works as a stress relieving agent (Höferl et al. 2006). In contrast, in catfish the (S)-enantiomer acts as a sedative (Heldwein et al. 2014). Here, we demonstrate that a racemic mixture of the two enantiomers of linalool enables live imaging of *Hydra* under various mounting and lighting conditions, including the acquisition of fluorescence time-lapse movies and multichannel z-stacks at high magnification. Linalool is fast acting – a 1mM solution of linalool anesthetizes an animal within 10 min of exposure, with recovery occurring in approximately the same time after removal from the solution. Because anesthesia using linalool is reversible, the same animal can be imaged consecutively over the course of days, enabling dynamic studies of long-term processes such as head regeneration and budding. Furthermore, linalool facilitates the rapid execution of precise sample manipulations such as tissue excisions and grafting. Linalool has been reported to be a cytostatic agent in cancer cells *in vitro* (Rodenak-Kladniew et al. 2018); therefore, we also investigated this possibility in *Hydra*. We found no significant effects of prolonged (3-day) continuous linalool exposure on budding rates, mitotic activity, or cell viability. In contrast, in amputated animals 3-day continuous exposure to linalool suppressed both head and foot regeneration but could be rescued upon removal of the anesthetic. Thus, linalool may also be a useful tool for manipulating regeneration dynamics. In conclusion, we find that linalool significantly outperforms other currently used anesthetics and enables *in vivo* manipulations and live imaging of *Hydra* with precision and ease of use.

## Materials and Methods

### *Hydra* strains and culture

We used the *Hydra vulgaris* AEP strain (Martin et al. 1997; Technau et al. 2003) and various transgenic lines derived from this strain: GCaMP6s, expressing the calcium sensor GCaMP6s in interstitial cells (Dupre & Yuste 2017); Wnt, expressing GFP under control of the Wnt3 promoter (Hobmayer et al. 2000); HyBra, expressing GFP under control of the HyBra2 promoter (Glauber et al. 2013); “Watermelon” (WM) animals (Glauber et al. 2013) expressing GFP in the ectoderm and DsRed2 in the endoderm with both genes under control of an actin gene promoter; and a line originating from a single animal that was obtained by recombining AEP ectoderm and watermelon endoderm following tissue separation (Cochet-Escartin et al. 2017) and named “Frank” by the undergraduate student who created it. The Frank line has unlabeled ectoderm and DsRed2-expressing endoderm. Nerve-free animals were generated by heat-shocking *H. vulgaris* strain A10. A10 is a chimeric animal consisting of *H. vulgaris* (formerly *Hydra magnipapillata* strain 105) epithelial cells and sf-1 interstitial cells (Shimizu et al. 2004a).

*Hydra* strains were maintained in mass cultures in *Hydra* medium (HM) composed of 1 mM CaCl_2_ (Spectrum Chemical, New Brunswick, NJ), 0.1 mM MgCl_2_ (Sigma-Aldrich, St. Louis, MO), 0.03 mM KNO_3_ (Fisher Scientific, Waltham, MA), 0.5 mM NaHCO_3_ (Fisher Scientific), and 0.08 mM MgSO_4_ (Fisher Scientific) prepared with MilliQ water, with a pH between 7 and 7.3. Cultures were maintained at 18°C in the dark in a Panasonic incubator (Panasonic MIR- 554, Tokyo, Japan). The cultures were fed 2-3x/week with *Artemia* nauplii from the San Francisco Bay or from the Great Salt Lake (Brine Shrimp Direct, Ogden, UT). Animals were cleaned daily using standard cleaning procedures (Lenhoff & Brown 1970). Asexual, non-budding polyps starved for at least 24 h were used for experiments unless stated otherwise.

### Generation of nerve-free *Hydra*

To generate nerve-free *Hydra*, A10 polyps were heat-shocked in an incubator (Fisher Scientific 615F) at 28-29°C in the dark for 72h and then moved back into the 18°C incubator (Sugiyama & Fujisawa 1978; Fujisawa 2003; Shimizu et al. 2004b). All nerve-free animals were subsequently force-fed and “burped” as described previously (Tran et al. 2017) for three to four weeks, in which time they lost nematocytes, as well as feeding and mouth opening behaviors.

### Preparing anesthetic solutions

Stock solutions were made in HM at concentrations of 1 mM linalool (Sigma-Aldrich), 0.04% heptanol (Acros Organics, Fisher Scientific), 2% urethane (Sigma-Aldrich), or 0.1% chloretone hemihydrate (Sigma-Aldrich). Linalool and heptanol were prepared fresh daily and stored at room temperature. Urethane and chloretone solutions were stored at 4°C for a few days and pre-warmed to room temperature before usage. Anesthetic solutions were prepared at room temperature, except for chloretone, which was prepared with slight heating.

### Linalool viability assay

24 h starved polyps were incubated in 6-well plates (Genesee Scientific, El Cajon, CA), 8 or 10 animals per well in 2 mL of different concentrations of linalool (0-10 mM) at room temperature for 3 h. The fraction of live animals was scored at the end of the assay. To obtain the LC50 value, the fraction of dead animals (1- fraction of live animals) was plotted against the linalool concentration. The data were fitted to the Hill equation as in (Hagstrom et al. 2015):

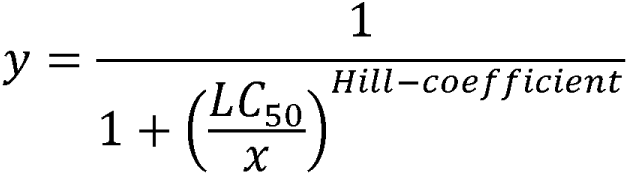

Here, y is the fraction of dead animals and x is the concentration of linalool in millimolar. The fit was generated using the curve fitting application in MATLAB (MathWorks, Natick, MA, USA)

### Characterizing short term efficacy of anesthetics

1-5 intact *Hydra* polyps were incubated per well in a flat bottom 6-well plate (Eppendorf, Hamburg, Germany) filled with 8mL of HM or respective anesthesia. If more than 2 polyps were used, 40 µm or 100 µm Falcon cell strainers (Fisher Scientific) were used in the well to allow for quicker transfer of the animals from HM to anesthesia and vice versa. In some experiments, all wells were imaged simultaneously and polyps were stained with neutral red (1:400,000 w/v; Fisher Scientific) in HM for 90 s at room temperature prior to the experiment to enhance contrast during imaging. The 6-well plate was imaged from the top using a Basler A601f-2 camera attached to a 25 mm TV lens C22525KP with adjustable focal length (Pentax, Tokyo, Japan) for 1 h at 1 fps using Basler pylon camera software. Lighting was provided by a model A4S light box (ME456, Amazon, Seattle, WA). After 1 h, the cell strainers were moved to a new well and 8 mL of HM were added to each well. Following this, the plate was imaged for 2 h. In other experiments, individual wells were imaged on a stereo microscope using a Flea-3 camera (FLIR Integrated Imaging Solutions Inc, Wilsonville, OR) controlled by a custom MATLAB script. To obtain representative images at higher magnification, anesthetized *Hydra* were imaged in a 35 mm tissue culture dish with a Leica MZ16FA microscope equipped with a SPOT RT3 camera (SPOT Imaging, Sterling Heights, Michigan), using the SPOT 5.1 software (SPOT Imaging) at 15min and at 60 min exposure.

A range of sublethal linalool concentrations (0 mM, 0.1 mM, 0.25 mM, 0.5 mM, 0.75 mM,1 mM) were tested. Working concentrations for other anesthetics were 2% urethane, 0.04% heptanol, or 0.1% chloretone, with induction imaged for at least 20 min and recovery for at least 30 min. At least 10 animals were assayed for each condition, in at least 3 technical replicates. Time of induction of anesthesia was considered to be the time at which the animal stopped extending further, and time of recovery was considered to be the timing of the first contraction burst observed after returning the polyps to HM. Due to the complex behavior of *Hydra* and the subjectivity of these measures, calculated times for induction and recovery should be considered estimates rather than conclusive values.

### Body column length of *Hydra* in anesthetics

24 h starved polyps were imaged for 10 min in HM to observe both extended and contracted states of the moving polyp to calculate an average body length ((max+min)/2). The polyps were then transferred to 1 mM linalool, 2% urethane, 0.04% heptanol, or 0.1% chloretone and imaged for an additional 20 min. We averaged the minimum and maximum body lengths of *Hydra* in the last 10 min of recording in each anesthetic. Average body length in the anesthetic was divided by the average in HM to find the % body length for each anesthetic to determine whether the polyps were hyperextended (>100%) or contracted (<100%) compared to their “normal” length. Because *Hydra* doesn’t have a fixed body shape or length due to constant extension and contraction, this normal length is somewhat arbitrary; however, it nevertheless allows us to compare the effects of the various anesthetics.

### Feeding and pinch responses in linalool

24 h starved polyps were incubated in 1 mM linalool for 10 min in a 60 mm tissue culture dish (VWR International, Radnor, PA). Each animal was pinched using a pair of Dumont No. 5 forceps (Fine Surgical Tools, Foster City, CA) to determine presence or absence of a contractile response while in the linalool solution. To assay whether animals exhibited a feeding response in linalool, 4-day starved polyps were first incubated for 10 min in 1 mM linalool. The anesthetized animals were transferred to a stereo microscope, and video recording with a Flea-3 camera controlled by a custom MATLAB script was started. Brine shrimp were added, taking care to only add a small amount of HM when transferring the shrimp, and recording was continued for 30 min. 4-day starved control animals in HM were imaged in the same way.

### Cross sections and “zebra grafts”

48-72h starved Wnt and Frank polyps were used. Polyps were placed in the lids of 35 mm dishes in either HM or 1 mM linalool for at least 10 min. Rings of tissue were excised from the body column using a scalpel 10 blade. The rings were strung onto glass needles pulled from microcapillaries (World Precision Instruments, Sarasota, FL) using a P-1000 micropipette puller (Sutter Instrument, Novato, CA) and imaged with a Leica MZ16FA microscope equipped with a SPOT RT3 camera, using the SPOT 5.1 software. “Zebra grafts” (n=2 per condition) were created using WM and Frank animals. The animals were placed in a 100 mm petri dish (Spectrum Scientifics, Philadelphia, PA) filled with either HM or 1 mM linalool in HM. A small piece of filter paper (2×2 mm) was cut and threaded onto a size 00 enameled insect pin (Austerlitz, Carolina Biological) and the pin placed into the dish. One animal was decapitated, and the head threaded onto the pin mouth first using forceps such that the cut edge of the tissue faced towards the point of the pin. The second animal was then decapitated and the head discarded. A ring of tissue was cut as thinly as possible from the body column of the second animal and threaded onto the pin, followed by a ring from the first animal. Alternating rings of tissue were cut and placed on the pin until the body columns of both animals were used up, at which point one of the feet was threaded onto the pin to complete the chimera. A second piece of filter paper was threaded onto the pin, and forceps used to gently move the two pieces of paper together in order to force all the rings into contact with each other. These chimeras were allowed to heal on the pins for 2 h, then gently pushed off the pins with forceps, transferred to clean 35 mm dishes full of HM, and allowed to further heal overnight before imaging.

Grafting of heads into the body column was accomplished using WM and unlabeled animals using an approach similar to the insect pin method described above. The WM animal was decapitated, and a slit cut in the side of the unlabeled animal. The pin was passed through the WM head hypostome first, then through the wound in the unlabeled polyp and out through the body wall on the other side. Care was taken when positioning the filter paper pieces to avoid pushing the donor head into the body cavity. Animals were allowed to heal for 2 h, then removed from the pins and placed in dishes of clean HM to heal overnight before imaging. Grafting of head organizers into the body column was accomplished without pins. Head organizers were obtained by anesthetizing a WM animal in linalool, removing the head, then excising the tentacle bases to leave only a small fragment of tissue containing the tip of the hypostome. A small slit was cut in the body column of an unlabeled animal, and forceps used to place the hypostome piece into the slit. Animals were allowed to heal for 2 h before transfer to dishes of clean HM. Successful grafts were imaged every 24 h to determine whether an ectopic body axis was induced.

### Fluorescence imaging in 1mM linalool and in other anesthetics

All imaging was done using an Olympus IX81 inverted microscope (Olympus Corporation, Tokyo, Japan) with an ORCA-ER camera (Hamamatsu Photonics, Hamamatsu, Japan). Slidebook software version 5.0 (Intelligent Imaging Innovations, Denver, CO) was used to interface with the microscope and acquire z-stacks and time-lapse images. Anesthesia incubations were performed as described earlier. *Hydra* expressing GCaMP6s and WM *Hydra* were used for fluorescence imaging. For low magnification single channel imaging, an animal was allowed to move freely in a drop of either HM or 1mM linalool on a 40 mm × 24 mm glass coverslip (Fisher Scientific) and was imaged in the GFP channel with a 50ms exposure using a 4x UPLFLN objective (Olympus). Images were recorded every 100ms for 10s to obtain a time lapse movie. Rigid body correction of z-stacks was accomplished using a previously described algorithm (Thévenaz et al. 1998). For high-magnification single channel imaging, animals were mounted in tunnel slides prepared as described in (Carter et al. 2016). Neurons in the body column of GcaMP6s animals were imaged by taking z-stacks of the tissue in the GFP channel (500ms exposure; z-step size of 0.25 µm), using a 60x oil immersion objective.

For low-magnification multi-channel imaging, WM animals were incubated in Hoechst 33342 (Thermo-Fisher Scientific) diluted 1:500 in 1mM linalool for 15 minutes in the dark. The animals were then decapitated and the hypostome mounted in a tunnel slide. Z-stacks were taken in DAPI, GFP and RFP channels with a step size of 2.99 µm using a 10x objective. For high-magnification multi-channel imaging, RWM animals were first incubated in SYTO 60 red fluorescent nucleic acid stain (Invitrogen) diluted to 10 µM in HM for 1 h at room temperature in the dark. 2 quick washes in 1mL HM followed, as well as a 15 min incubation in the dark at room temperature in 1:250 Hoechst 33342 diluted in 1mM linalool. Body columns of the animals were imaged in the DAPI, RFP and DRAQ5 channels with a 60x oil immersion objective. For high magnification two-channel imaging, Hoechst 33342 (Thermo-Fisher Scientific) was diluted 1:500 in 1 mL of the respective anesthetic solution and WM animals were incubated for 15 min at room temperature in the dark. Individuals were mounted on tunnel slides and imaged.

### Regeneration and budding assays

Polyps were decapitated with a scalpel just below the tentacle ring for head regeneration and above the budding zone for foot regeneration assays. In one experiment, the decapitated animals were placed in 600 µL of 0 mM (control), 0.1 mM, 0.25 mM, 0.5 mM, 0.75 mM, or 1 mM linalool in HM. Head regeneration was scored by the appearance of the first tentacle on a decapitated animal. 8 animals were kept at each concentration in a 48-well plate (Eppendorf) and imaged in brightfield at 4x with an Invitrogen EVOS Fl Auto 2 (Thermo-Fisher Scientific, Waltham, MA). Head regeneration was scored every 12 h for 72 h. The lid of the plate was removed for imaging and the solutions were changed every 24 h. In another experiment, decapitated polyps were placed individually into the wells of a 24-well plate (Eppendorf), filled either with 500 µl HM or 1 mM linalool. Polyps were imaged approximately every 12 h and the appearance of tentacles and hypostomes were scored. After approximately 3 days, polyps were transferred into a new 24-well plate containing 500 µl fresh HM and imaged a day after transfer. Foot regeneration experiments were conducted the same way, with animals scored for the appearance of a peduncle and for the ability to adhere to the substrate. For repeated imaging of head regeneration at high magnification, animals were anesthetized in 1 mM linalool for 10 min prior to imaging and returned to HM to recover afterwards. To facilitate removal from the slides, a layer of Scotch tape was placed over the double-sided tape during construction of tunnel slides. The increased space between coverslip and slide and ability to easily lift off the coverslip after imaging allowed recovery of the animal with minimal chance of injury.

Budding was assessed by selecting healthy animals with early buds at stages 3-4 on the previously described scale (Otto & Campbell 1977), and incubating them in well plates as described for regeneration assays. Animals were scored for development of tentacles on the bud and formation of further buds. Long-term imaging of budding was carried out in 35 mm glass bottomed dishes (MatTek, Ashland, MA). One animal was placed onto the glass surface at the bottom of the dish in 1 mM linalool, a coverslip was laid over the top to constrain the animal, and the dish was flooded with 1 mM linalool. Animals were imaged once per hour for 48 h using an Invitrogen EVOS FL Auto microscope.

### Cell viability assay

Polyps were incubated for 30 min in 1 μg/mL propidium iodide in HM, washed twice in HM, then mounted on glass slides as described for live imaging of neurons. Slides were imaged on an Invitrogen EVOS FL Auto microscope in the red fluorescence channel using the Invitrogen EVOS FL Auto Imaging System software. Labeled cells were counted in the body column only and reported as number of labeled cells per animal. As a positive control, polyps were incubated in 0.04% colchicine (Acros Organics) in HM to induce cell death (Cikala et al. 1999). Animals were incubated in colchicine for a full 24 h rather than 8 h incubation followed by 16 h recovery as described.

### Mitotic index assay

Polyps were incubated in HM or 1 mM linalool for 72 hours in 60 mm cell culture dishes at a density of 1 polyp/mL. Polyps were not fed during the experiment, but the medium was changed daily. At the end of the 72 h, one or two cross sectional segments were cut from the body column of each polyp near the head. The samples were placed on glass slides for a wet mount antibody stain. Humid chambers for staining were constructed by lining covered 100 mm Petri dishes (Spectrum Scientific) with wet paper towels and placing the slides inside the dishes. A well was created in the center of each glass slide by layering two pieces of double-sided tape across both short sides of the slide with one piece of tape running on both long edges of the slide. The samples were placed in a drop of medium on the slide. All steps were performed at room temperature unless otherwise noted. The samples were fixed in 20 µL 4% paraformaldehyde (Sigma-Aldrich) in HM for 15 min. The samples were washed three times with 20 µL 1x PBS, followed by a 15 min permeabilization with 20 µL 0.5% PBSTx (0.5% Triton-X in 1x PBS). They were then incubated for 3.5 h in 20 µL blocking solution (1% FBS, 0.1% DMSO in 1x PBS) and placed overnight (16h) at 4°C in 30 µL anti-phospho-histone H3 (Ser10) primary antibody (Millipore Sigma, Burlington, MA) diluted 1:100 in blocking solution. On the second day, samples were washed quickly 3x with 40 µL 1x PBS, followed by four 25-35 minute washes of 20 µL 0.3% PBSTx. The samples were then incubated in a 1:1000 dilution of Alexa 546 rabbit IgG secondary antibody (Thermo-Fisher Scientific) for 5 h, followed by three quick and two 10 min washes of 0.3% PBSTx. To stain nuclei, the samples were incubated in DRAQ5 (Thermo-Fisher Scientific) diluted to 5 µM in 1x PBS for 15 min and then washed three times with 1x PBS. The 1x PBS was replaced with a 1:1 solution of glycerol and HM. Finally, a cover slip was placed over the samples and nail polish was used to seal the slides. Z-stacks of the cross-sections were imaged using a Leica high-resonance scanning SP5 confocal microscope with a 20x C-Apochromat 1.2 W objective.

To calculate mitotic indices, the number of Alexa 546 stained nuclei was counted for each cross section, divided by the number of nuclei stained by DRAQ5 and multiplied by 100 to obtain a percentage. Counting of Alexa 546 and DRAQ5 stained nuclei was done using Fiji (Schindelin et al. 2012). For the z-stack corresponding to each color channel, a maximum intensity z-projection was taken and binarized. The projection was then segmented using the water-shedding tool. The number of particles was counted using the *Analyze Particles* tool, with a size range of 10- infinity µm^2^. For Alexa 546 color channel stacks, an additional thresholding step was used before binarizing the image.

## Results

### Linalool is a fast acting and reversible anesthetic

Intact polyps in HM continuously exhibit body shape changes and tentacle movements, contracting, extending, and bending, which greatly complicates *in vivo* manipulations and imaging. In contrast, animals incubated in 1mM linalool for 10 min appear relaxed, with tentacles splayed out and the mouth assuming a conical shape (Fig.1 A).

**Figure 1.**
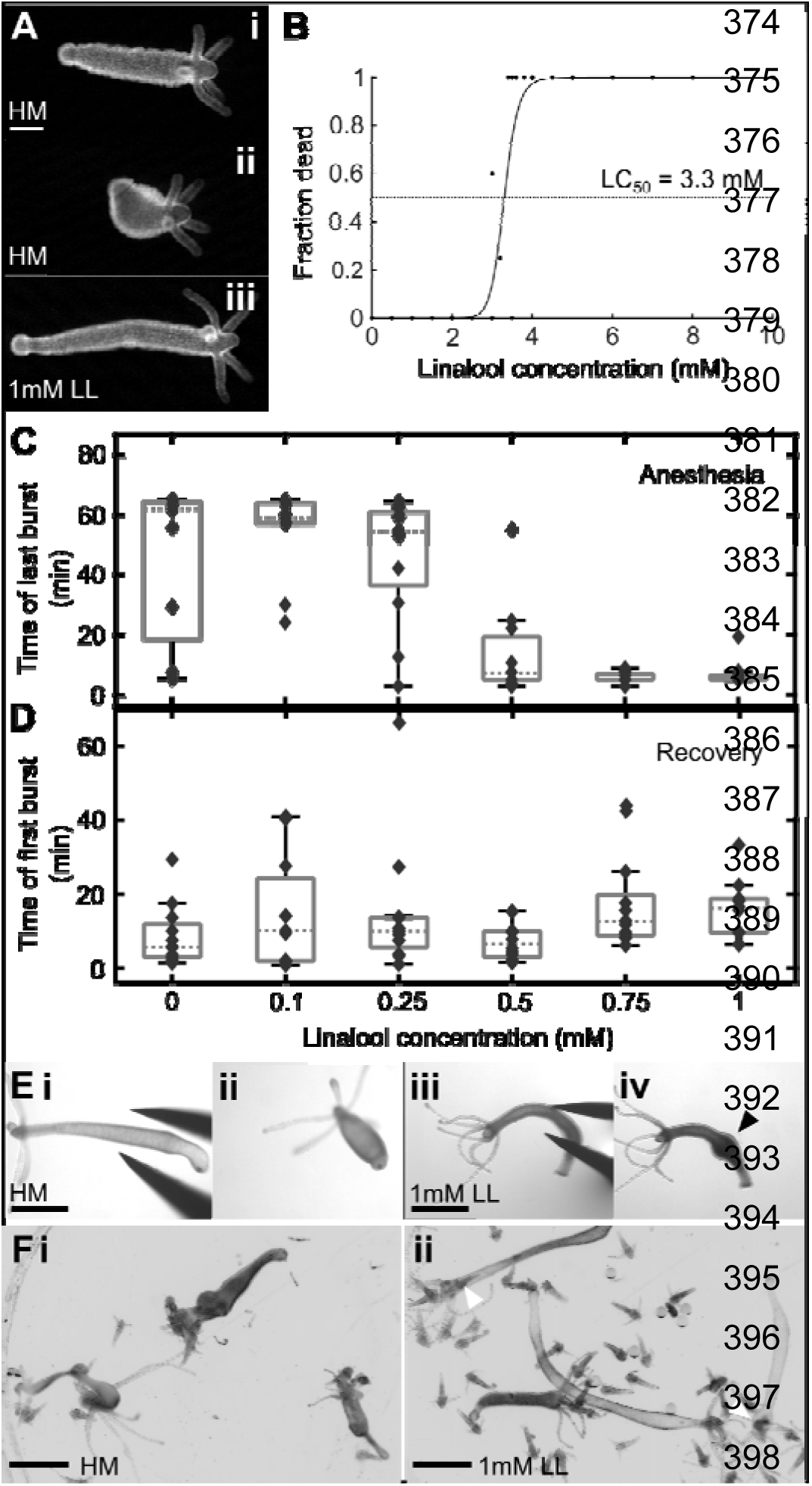
Linalool as an anesthetic. A. Representative images of *Hydra* polyps before (i,extended, ii, contracted) and after (iii) incubation in 1 mM linalool (abbreviated to LL). Scale bar: 200 µm. B. 3 hour incubation in linalool concentrations exceeding 2 mM causes lethality. C. Box plot showing time of last observed contraction burst during 60 min incubation in linalool concentrations up to 1 mM. D. Box plot showing time of first observed contraction burst during 60 min recovery in HM following 1h of anesthesia in linalool. E. Pinch response. i. *Hydra* polyp in HM. ii. Polyp in HM shows a contractile response to pinching. iii. *Hydra* polyp incubated in 1 mM linalool for 10 min. iv. Anesthetized polyp shows only local swelling after pinch, indicated by black arrowhead. F. 30 min feeding response in 4-day starved polyp. i. *Hydra* polyps in HM readily capture and consume multiple shrimp. ii. *Hydra* polyps incubated in linalool for 10 min prior to introduction of shrimp have a much reduced reaction, and only rarely ingest shrimp. White arrowheads indicate shrimp inside polyps. Scale bars for E, F: 1 mm.

We investigated the effect of various linalool concentrations on animal health within 3 hours of incubation (Fig.1B) and found that concentrations exceeding 2 mM caused negative health effects on the animals, such as an abnormal body shape, contracted tentacles, and partial disintegration. Death was observed at concentrations of 3 mM and beyond following 3h exposure. We determined the LC_50_ to be 3.3 mM using the same approach as in (Hagstrom et al. 2015). We then empirically determined the optimal working concentration for linalool by measuring and comparing induction and recovery times for different sublethal concentrations.

No negative health effects were observed at or below 1mM linalool. Induction time of anesthesia decreased with increasing concentration of linalool to about 10 min at 1mM (Fig 1C) while recovery time remained constant (Fig. 1D) at 10-20 min for all concentrations tested. Therefore, we determined that the highest tolerated dose, 1mM, was the best concentration to use in experiments. Polyps incubated in 1mM linalool for at least 10 min no longer exhibit the “pinch response”, a global longitudinal contraction, that is observed in HM upon gently squeezing the body column with forceps (Fig.1 E i). Polyps in 1 mM linalool swelled at the site of pinching but did not contract (Fig.1E ii). Thus, linalool causes *Hydra* to lose both spontaneous body column contractions and mechanically induced ones. Mechanically induced body column contractions are known to be mediated by the ectodermal epithelial layer, and nerve-free animals retain their pinch response despite lacking spontaneous contraction behaviors (Takaku et al. 2015). The loss of both upon treatment with linalool suggests that linalool affects both the neuronal and the epithelial cells. However, 1mM linalool does not completely paralyze the animal, as we observed that a few anesthetized individuals were able to capture and ingest shrimp, although very inefficiently compared to controls (Fig.1 F).

### 1 mM linalool enables precise tissue manipulations

Recent studies have shown that the regeneration outcome in *Hydra* could be influenced by the geometry of tissue pieces excised from the body column (Livshits et al. 2017). To test whether linalool allowed for improved precision of cuts and thus would be a useful tool for such studies, we compared the excision of tissue rings from animals incubated in HM with those incubated in 1 mM linalool. When sectioning animals to obtain pieces of body column tissue, the application of linalool does not drastically improve minimum possible slice thickness. However, it significantly reduces the working time required, from approx. 3 min to 30 s per animal (Fig. 2A). This is due to the suppression of the animal’s natural contractile response to touch, removing the need to wait for the polyp to extend following each cut.

**Figure 2.**
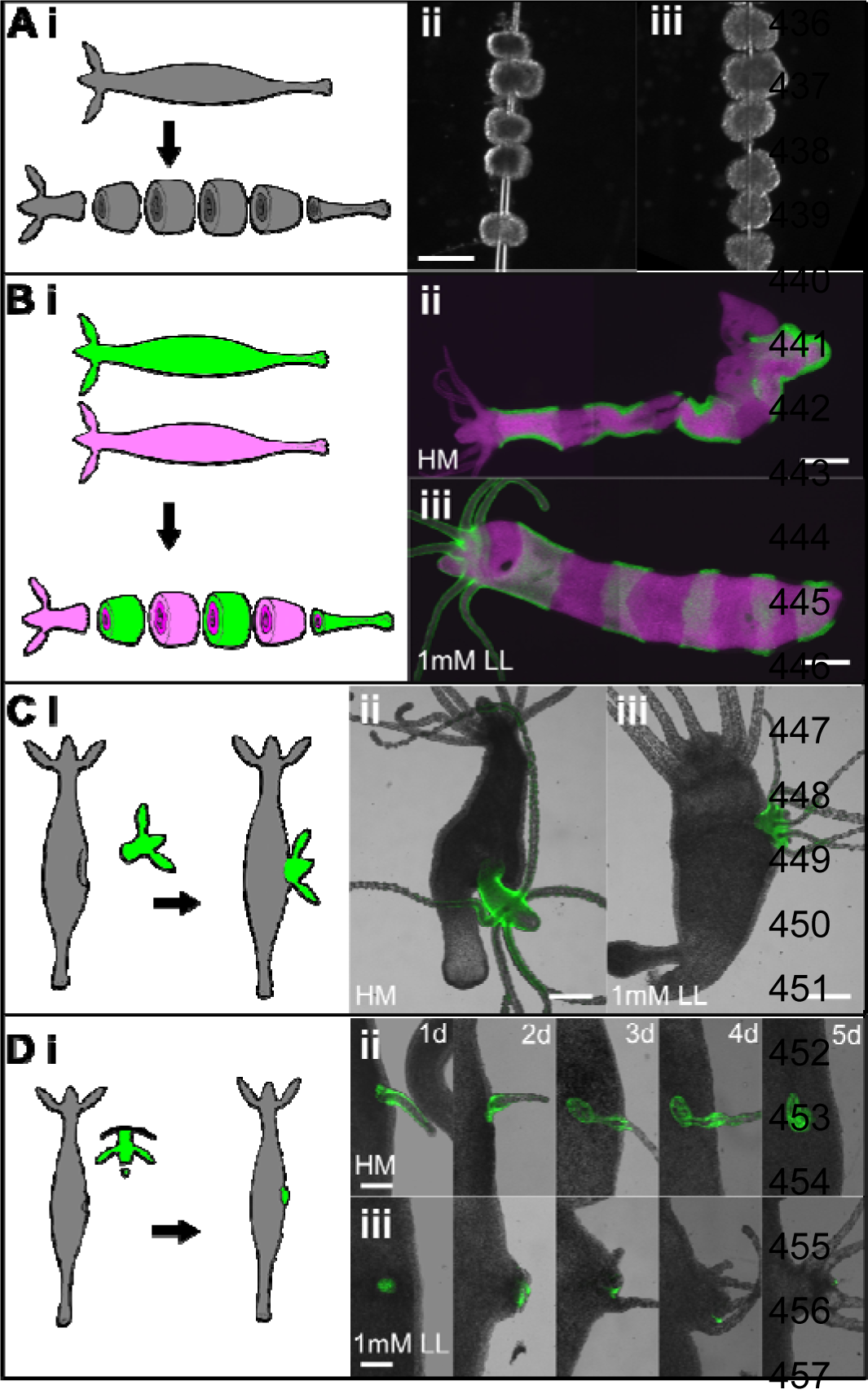
Linalool improves outcomes of surgical manipulations in *Hydra*. A. Sectioning of body column. i. Experimental schematic. ii. Sections cut in HM. Slices have average thickness (197 ±7) µm (mean± std) and took about 2 min 51s per animal averaged over 3 animals. iii. Sections cut in linalool. Slice thickness 178 ± 4µm, average time 36s over 3 animals. Scale bar: 400 µm. B. “Zebra grafting”. i. Experimental schematic. ii. Representative animal grafted and healed in HM. iii. Representative animal grafted and healed in linalool. Scale bars: 400 µm. C. Head transplantation into gastric region. i. Experimental schematic. ii. Representative animal grafted and healed in HM. iii. Representative animal grafted and healed in 1mM linalool. Scale bars: 400 µm. D. Head organizer transplantation into gastric region. i. Experimental schematic. ii. Animal grafted in HM imaged over 5 days. iii. Animal grafted in 1 mM linalool imaged over 5 days. Scale bars: 200 µm.

The improvements possible using linalool become more readily apparent in grafting experiments. A “zebra graft” to create a chimeric animal consisting of bands of differently labeled tissue produced a significantly better result when linalool was employed (Fig. 2B). Linalool incubation roughly halved the time required to cut the rings of tissue and thread them onto the needle, but the true benefit is in immobilization of the tissue during the initial 2 hour healing step on the needles. Grafts looked similar immediately after their creation but animals grafted in HM had abnormal morphology immediately apparent on removal from the needles (Fig. 2B ii, iii). This is likely due to tissue movement causing the cut edges of the pieces to become misaligned while on the needle, thus preventing the segments from healing smoothly together as described previously (Shimizu & Sawada 1987). A similar effect was observed when grafting heads onto body columns (n=3 per condition), as previously described (Rand et al. 1926). Linalool allowed more precise decapitation of the donor animal, reducing the amount of extraneous body column tissue, and guaranteed better positioning of the graft on the recipient animal. Grafts carried out in HM tend to have the donor head protruding at an angle, again due to misalignment of the cut surfaces during healing (Fig. 2C). Finally, linalool improves overall outcome in hypostome grafts carried out as previously described (Broun & Bode 2002). Grafts in HM incorporated tissue that formed only an ectopic tentacle before being resorbed (Fig. 2D ii) or resulted in an entire head formed from donor tissue (data not shown). Grafts in linalool resulted in formation of an ectopic body axis from recipient tissue, with donor tissue limited to a small part of the new head (Fig. 2 D iii), as previously described (Browne 1909; Broun & Bode 2002).

### Incubation in 1 mM linalool enables high-quality fluorescence short-term imaging

To test whether the immobilization in 1 mM linalool was sufficient to allow for *in vivo* fluorescence imaging, we imaged animals incubated in 1mM linalool under various conditions and compared the results to those obtained from imaging animals in HM. First, we used single channel fluorescent imaging using polyps expressing GCaMP6s in the interstitial cell lineage (Dupre & Yuste 2017), because this transgenic line allows for the visualization of individual neurons and subcellular processes such as dendrites. Because GCaMP6s animals were originally developed to study neuronal control of behavior in *Hydra*, we imaged unconstrained animals at low magnification (Fig. 3A, Movie S1). Unconstrained animals in HM moved significantly during the 10 s acquisition, as shown by a maximum intensity projection of the time series (Fig.3 A ii). In contrast, polyps incubated in 1 mM linalool for at least 10 min only exhibited drift (Fig.3 A iv, v), which can be corrected for with standard post-processing methods (Fig. 3A vi), whereas these methods do not correct for the motion observed in the control, because the animal exhibits non-linear body shape changes (Fig. 3A iii).

**Figure 3.**
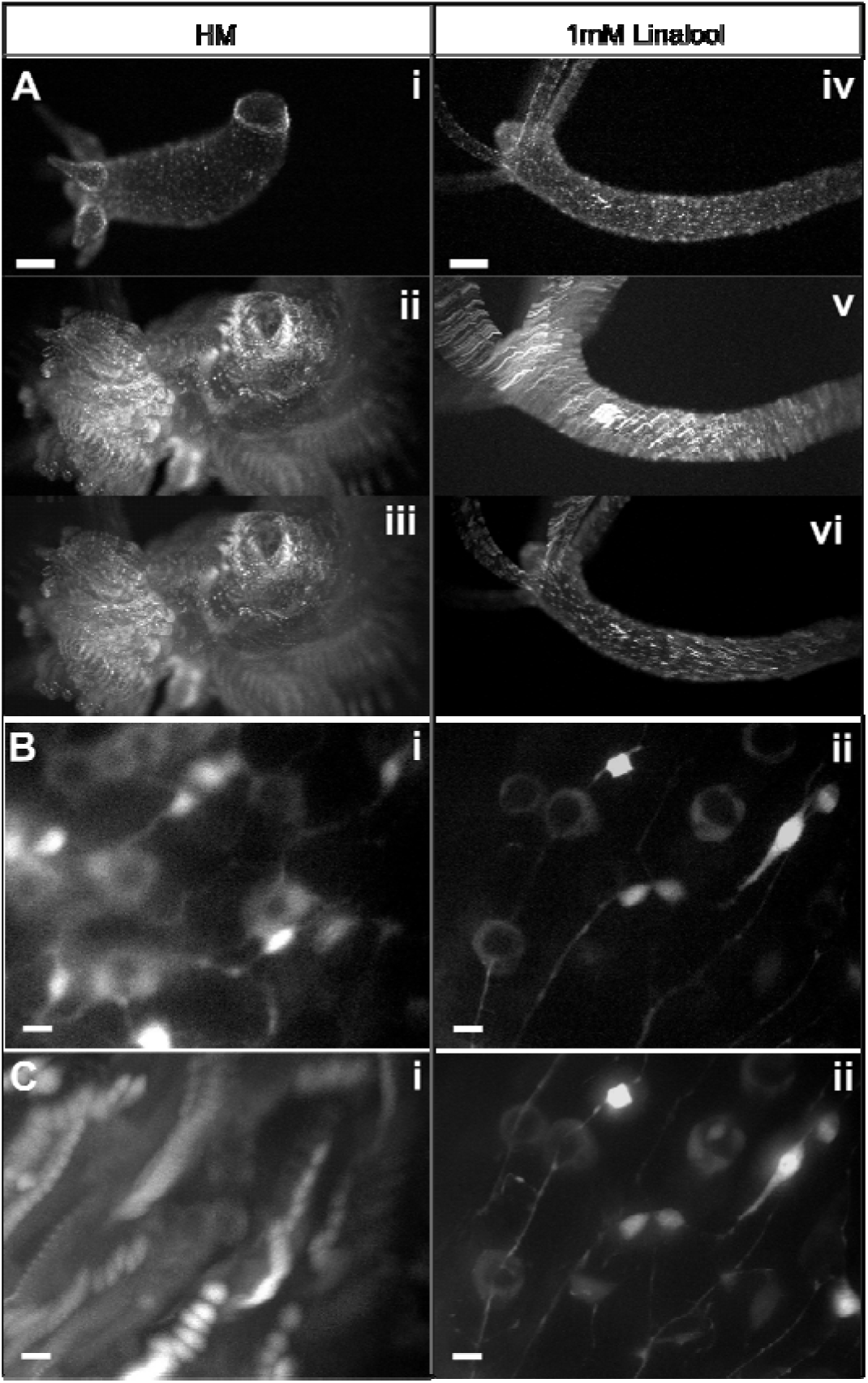
Live imaging in linalool. A. Unconstrained GCaMP6s *Hydra* imaged at low magnification. i. single image in HM. ii. Maximum intensity Z-projection of a 10 s video in HM. iii. Rigid body correction of HM video projection. iv. Single image in 1 mM linalool. v. Maximum intensity Z-projection of 10 s video in linalool. vi. Rigid body correction of linalool video projection. Scale bars: 200 µm. B. Single slice from a z-stack of a GcaMP6s animal imaged at 60x magnification with a resolution of 0.25 µm along the z-axis at a 200 ms exposure per slice using blue excitation in (i) HM and (ii) 1 mM linalool. C. Maximum intensity projection of high magnification z-stacks in (i) HM and (ii) 1 mM linalool. Scale bars: 10 µm.

We also acquired 20 µm thick z-stacks of the body columns of intact polyps mounted in tunnel slides (Carter et al. 2016) at high magnification (Fig. 3B, C). The image quality of individual slices was better when imaging anesthetized animals (Fig. 3B), but the difference in stability and thus image quality becomes most evident when comparing maximum intensity projections of the entire z-stack (Fig. 3C). The animal in linalool is sufficiently still to allow the resolution of subcellular features such as neuronal processes, whereas the animal in HM moves too much, making z-stacks impractical (Fig. 3C and Movie S2). As the tissue stretches and compresses anisotropically during those movements, it is not possible to correct this motion through post-processing.

Next, we tested the performance of 10 min incubation in 1mM linalool for the acquisition of multi-channel z-stacks at low (10x) and high magnification (60x). Control videos in HM were not attempted due to the unsatisfactory results obtained in a single channel (Fig. 3). By exposing animals to 1mM linalool in the presence of 2mM reduced glutathione, we were able to induce mouth opening (Fig.4A). The animal is sufficiently still to allow for simultaneous visualization of nuclei positions and cell boundaries. We also took 3-channel time-lapse movies of heads exposed to reduced glutathione below the activation threshold for opening to illustrate the overall stability that can be achieved using linalool, allowing for co-localization studies of dynamic processes (Movie S3). Finally, we tested whether animals were sufficiently immobile to obtain high quality z-stacks in multiple channels. While motion is not completely suppressed in linalool and extended exposure to short wavelength light causes the animal to escape the field of view, it is possible to achieve high quality multichannel imaging (Fig. 4B). Thus, linalool is a useful tool for *in vivo* co-localization studies. Notably, when testing live dyes, we found that the SYTO 60 red fluorescent nucleic acid stain is specific to nematocysts of all types in *Hydra*, determined by comparing morphology of stained structures to previous descriptions of nematocyst types (Engel et al. 2002).

**Figure 4.**
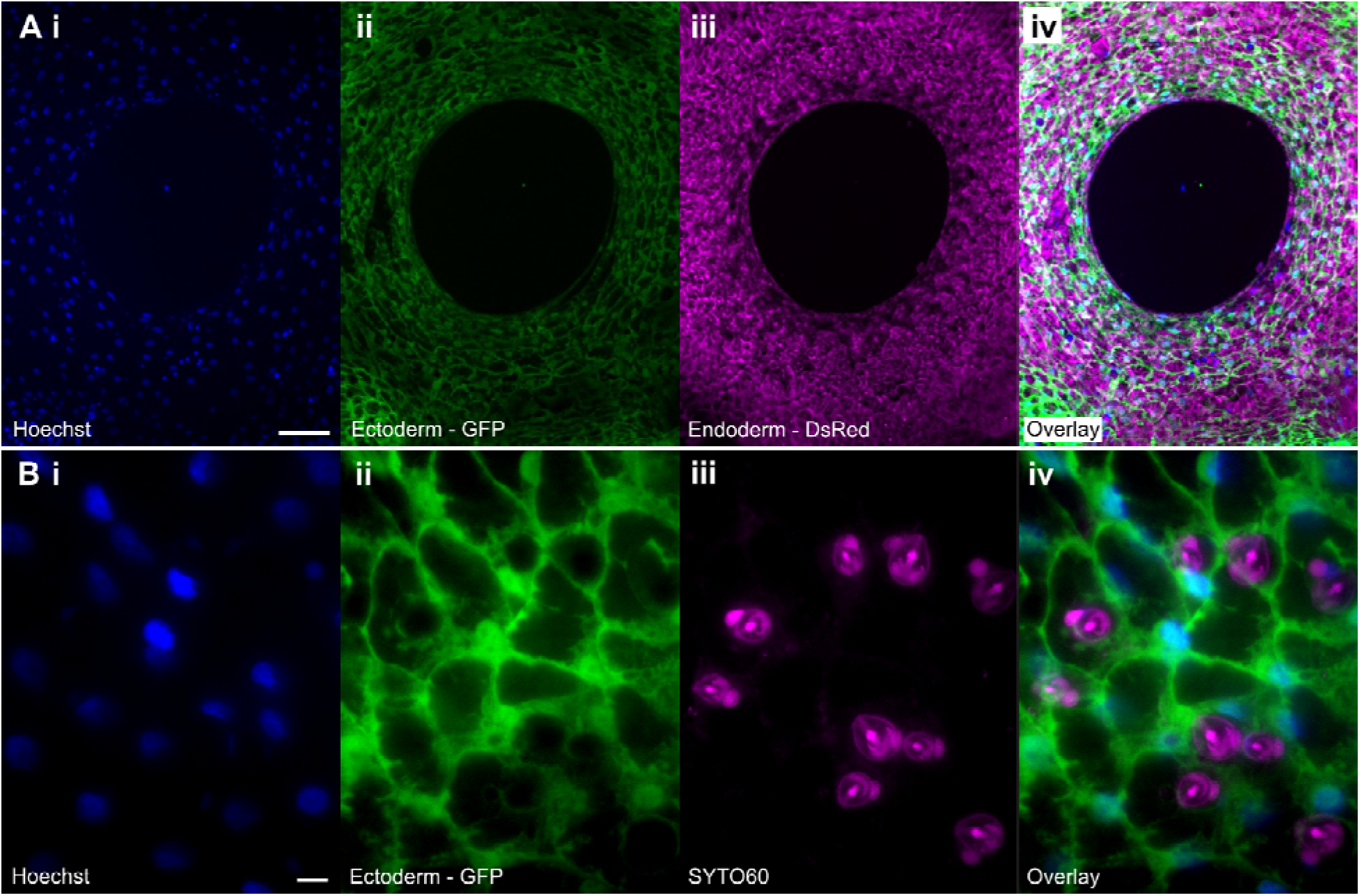
Linalool enables high resolution imaging in multiple channels. A. Low magnification maximum intensity projection of a z-stack acquired of an open *Hydra* mouth using i. Hoechst 33342, ii. Ectoderm – GFP, iii. Endoderm – DsRed2, iv. overlay. 5 µm slice thickness, 6 slices total. Scale bar: 100 µm. B. High magnification maximum intensity projection of a z-stack of the body column using i. Hoechst 33342, ii. Ectoderm – GFP, iii. Nematocysts – SYTO 60, iv. overlay. 0.25 µm z-step, 17 slices total. Scale bar: 10 µm.

### Repeated fluorescent short-term imaging

A major strength of linalool as a reversible anesthetic is the ability to repeatedly anesthetize and image the same animal over the course of days, thus allowing the acquisition of dynamic data of cellular processes in a single animal. To illustrate this capability, we decapitated transgenic HyBra2 promoter::GFP animals and allowed them to regenerate in HM. We imaged head regeneration over the course of 2 days, using repeated short-term 15 min incubations in 1mM linalool to acquire a total of 11 high resolution images of the same animal (Fig. 5A). When not imaged, the regenerating animals were returned to HM. In this way we were able to observe the development of the hypostome and tentacles and also to observe a gradual increase in GFP signal beginning at 24h. The same technique of repeated linalool exposure was used to image the tissue grafts in Figure 2.

**Figure 5.**
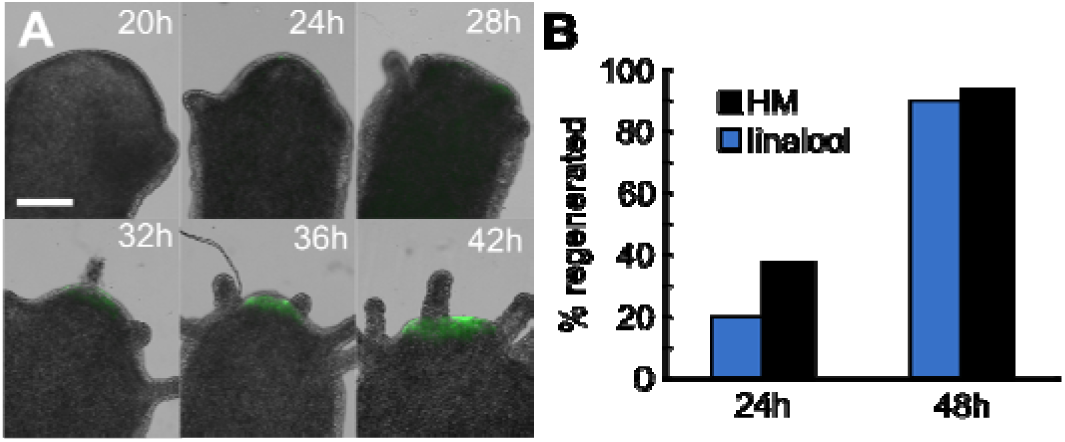
Linalool enables repeated high resolution imaging. A. Head regeneration in a transgenic HyBra2 promoter::GFP polyp imaged at high resolution every 4h from 12h to 48h. Subset of images shown. Scale bar: 0.5 mm B. Repeated anesthesia and recovery does not impact regeneration speed or outcome (n=10 animals HM, n=16 animals linalool, N=3 technical replicates). Differences between conditions not statistically significant at p=0.05 level.

We also confirmed that the timing and outcome of head regeneration in animals repeatedly anesthetized for imaging does not significantly differ from that observed in untreated controls. (Fig 5B). Thus, linalool is a valuable tool for repeated live imaging applications, which will be useful to study long term processes during regeneration and budding.

### Long-term applications of linalool

Due to the reported cytostatic effect of linalool on cancer cells in culture, (Rodenak-Kladniew et al. 2018) we investigated whether linalool has similar effects in *Hydra*. The cell cycle lengths in interstitial and epithelial cells are approximately 1 (Campbell & David 1974) and 3 days (David & Campbell 1972), respectively. Therefore, we continuously incubated intact polyps for 3 days in 1 mM linalool, exchanging the solution every 24 hours to account for volatility. We neither observed significant changes in mitosis (Fig 6A) nor in cell death (Fig. 6B) in the body column of intact polyps. Furthermore, budding seemed to occur normally, as verified using 3-day continuous time-lapse imaging (Fig. 6C, Fig. S1) and budding rates of polyps in 1 mM linalool were comparable to those of controls (Fig. 6D). Based on these results, we attempted to image head regeneration in 1 mM linalool. This would be advantageous compared to consecutive mounting and imaging sessions as it would minimize interaction with the sample and could be fully automated. However, we found that decapitated *Hydra* were unable to regenerate heads in 1 mM linalool when continuously exposed over the course of 3 days. Anesthetized body columns were observed to shed cells and assume a lollipop shape (Fig. 6E i, ii), and a few animals disintegrated completely. If removed from linalool after 3 d, however, the surviving animals recovered. Tentacle buds were observed as early as 1 day into recovery and all polyps had fully regenerated their heads after 3d of recovery. (Fig. 6E).

**Figure 6.**
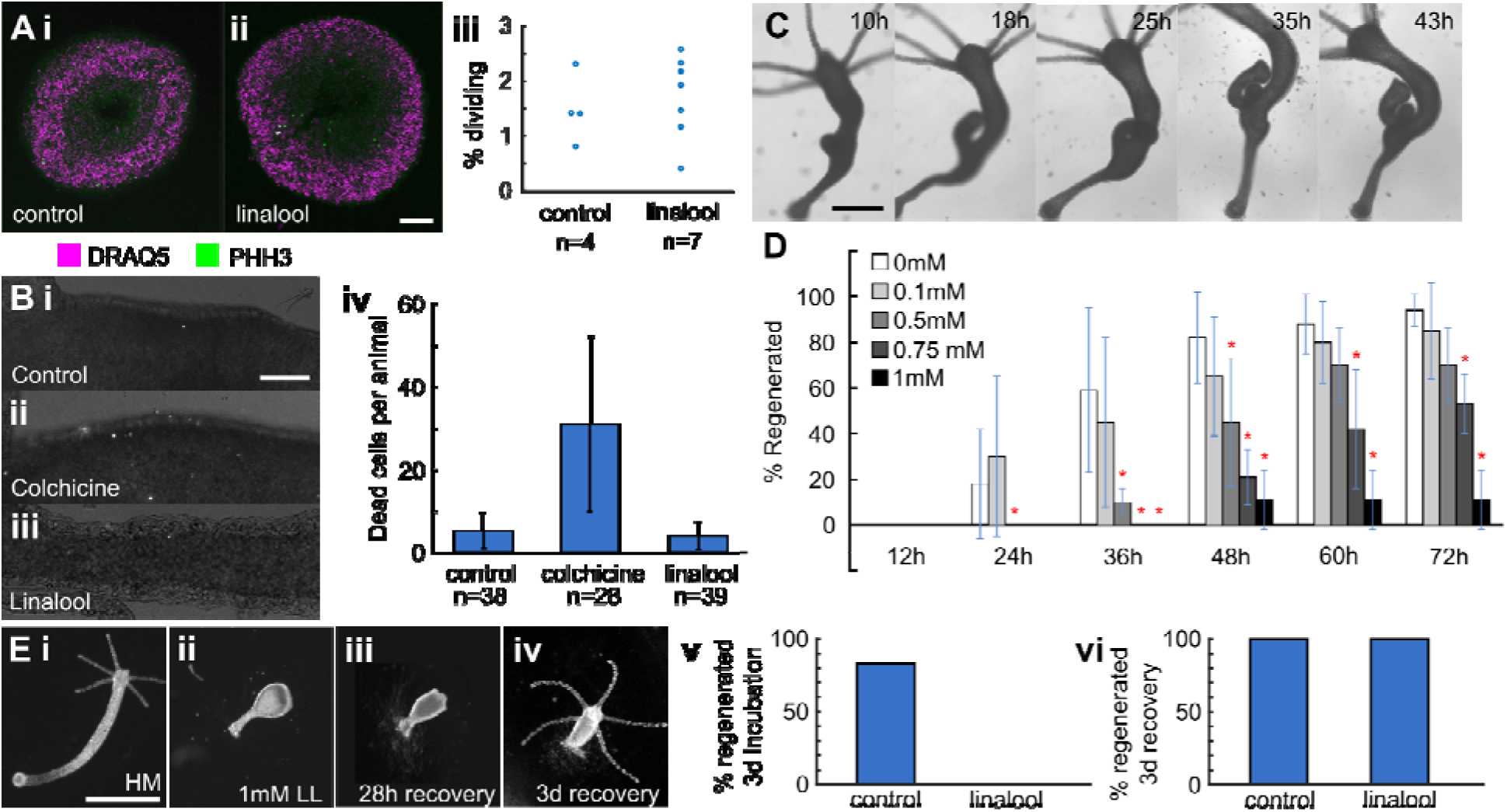
Effect of long-term continuous linalool exposure. A. 3 day incubation in 1 mM linalool does not impact rate of cell division. i. Representative image of body column sections from polyps incubated 3 days in HM. ii. Representative slice from polyps incubated 3 days in 1 mM linalool. Slices stained with DRAQ5 (nuclei) and anti-PH3 (phospho-histone H3, dividing cells). iii. Percentage of dividing cells in animals incubated 3 d in HM (control) or 1mM linalool. Mean ± standard deviation: control = 1.5 ± 0.6, linalool = 1.5 ± 0.7. Scale bar: 100 µm B. 3 day incubation in 1 mM linalool does not damage or kill cells. Representative images of polyps stained with propidium iodide after incubating for i. 3 days in HM, ii. 24 h in 0.04% colchicine, and iii. 3 days in 1 mM linalool. iv. Mean number of dead cells per animal. Error bars represent standard deviations. Scale bar: 100 µm C. Long term incubation in linalool does not impact budding. Representative images of a budding polyp continuously incubated and imaged in 1 mM linalool. Scale bar: 500 µm D. Long term incubation in linalool prevents head regeneration. Error bars represent standard deviations (0mM n=17, 0.1mM n=20, 0.5mM n=40, 0.75mM n=19, 1mM n=19; 3 technical replicates). Red asterisks indicate statistically significant difference from 0 mM as determined using Fisher’s Exact Test. E. Recovery in HM rescues the head regeneration defect. i. Polyp incubated in HM for 68 h after decapitation. ii. Polyp incubated in 1 mM linalool for 68 h after decapitation. iii. Decapitated polyp recovered for 28 h after 3 d in 1mM linalool, iv. Polyps recovered for 3 d after 3 d in 1mM linalool. Scale bar: 1 mm. v. Head regeneration is prevented by incubation for 3 d in 1 mM linalool, n=12. vi. Head regeneration in animals incubated in linalool for 3 d followed by recovery in HM for 3 d, n=6.

The negative effect of continuous linalool exposure on head regeneration was observed for concentrations as low as 0.5 mM (Fig. 6D). At 0.75 mM, 50% of the animals did not regenerate heads within 3 days. At 0.25 mM or lower, animals regenerated heads similarly to the control; however, these concentrations were ineffective in immobilizing animals for long-term imaging (data not shown). Foot regeneration was similarly suppressed under continuous 1 mM linalool exposure (Fig. S2), showing that the effects of linalool on regeneration are not specific to the head.

Finally, we tested whether the inhibition of regeneration is caused by an effect on the nervous system, as it had previously been suggested that the nervous system plays a role in head regeneration (Miljkovic-Licina et al. 2007). To this end, we generated nerve-free animals as described in Methods and assayed head regeneration in1 mM linalool. Surprisingly, nerve-free animals in 1 mM linalool regenerated similar to the controls maintained in HM. After 4 days of regeneration, 5/7 animals in HM and 3/7 in linalool showed tentacle buds. By 5 days this had increased to 7/7 in HM and 5/7 in linalool (Fig. S3). Furthermore, nerve-free animals in linalool never assumed the lollipop shape (Fig. 6E ii) that we observed in enervated polyps. Together, these data suggest that linalool disrupts regeneration by perturbing the function of either neurons or other cells in the interstitial lineage.

### Comparison of linalool to other commonly used anesthetics in *Hydra* research

Whenever one introduces a new tool, it is important to compare performance with existing methods and demonstrate that the advantages of the new tool are sufficient to make its adoption worthwhile. While anesthetics were and continue to be most frequently used to relax *Hydra* prior to fixation for histological and immunohistochemistry studies (Hausman & Burnett 1971; Benos et al. 1977; Münder et al. 2013; Buzgariu et al. 2018), the advent of modern molecular tools have brought with it an increased use for *in vivo* applications (Takahashi & Hamaue 2010; Badhiwala et al. 2018; Lommel et al. 2017). Table 1 provides an overview of the various anesthetics that have been reported in the literature for use in *Hydra* and examples of their respective applications.

**Table 1.**
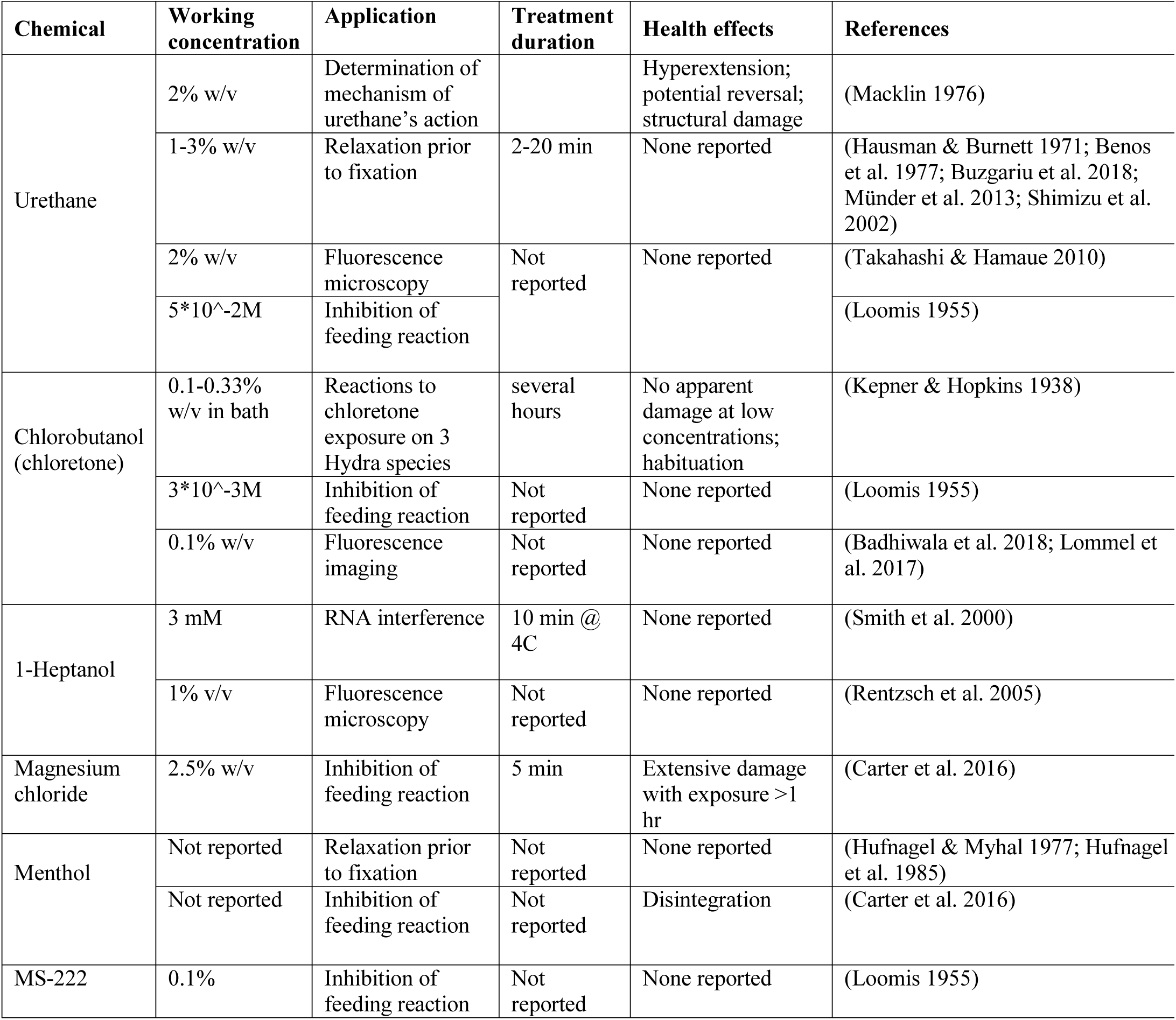
Summary of various anesthetics used to relax Hydra. This table is not a comprehensive summary of all *Hydra* studies that have employed anesthetics but provides an overview of examples spanning different chemicals and applications. To the best of our knowledge such a direct comparison has not previously been attempted and thus will be a useful resource for the field.

Based on our literature search, the most prominent *in vivo* application of the anesthetics was fluorescence imaging using urethane, heptanol, or chloretone. We therefore compared linalool to these anesthetics. To this end we studied whether there were any differences in morphology when *Hydra* polyps are exposed to the different substances. Although we observed variability among individual polyps exposed to the same anesthetic at a fixed concentration, both in terms of morphology and in terms of immobilization speed and strength, polyps assumed characteristic shapes upon exposure to the different chemicals (Fig. 7A). Following a 15 min exposure, *Hydra* polyps incubated in 1 mM linalool appear relaxed with tentacles splayed outwards and cone-shaped hypostomes (Fig 7A i). This morphology does not change significantly by 60 min. Animals incubated in 0.04% heptanol appear less extended at 15 min, with contracted conical tentacles. At 60 min the body columns are contracted and the stubby tentacles persist (Fig 7A ii). Exposure to 2% urethane causes animals to extend and become very thin at 15 min, though they become swollen while remaining extended by 60 min. (Fig 7A iii). 0.1% chloretone causes initial extension without the thinness seen in urethane, followed by the formation of swellings along the body column by 15 min and contraction of both body and tentacles by 60 min (Fig 7 A iv). To quantify these differences, we calculated average body length of individual animals after 10 min incubation in anesthetic as a percentage of their average length prior to anesthesia (see Methods). We found that linalool (median, (25^th^ percentile, 75^th^ percentile) = 103%, (87, 112)), heptanol (83%, (71, 93)) and urethane (96%, (88, 118) produce similar anesthetized lengths, while chloretone (133%, (125, 153)) shows a statistically significant increase in length and some hyperextended animals (Fig. S4A, D).

**Figure 7.**
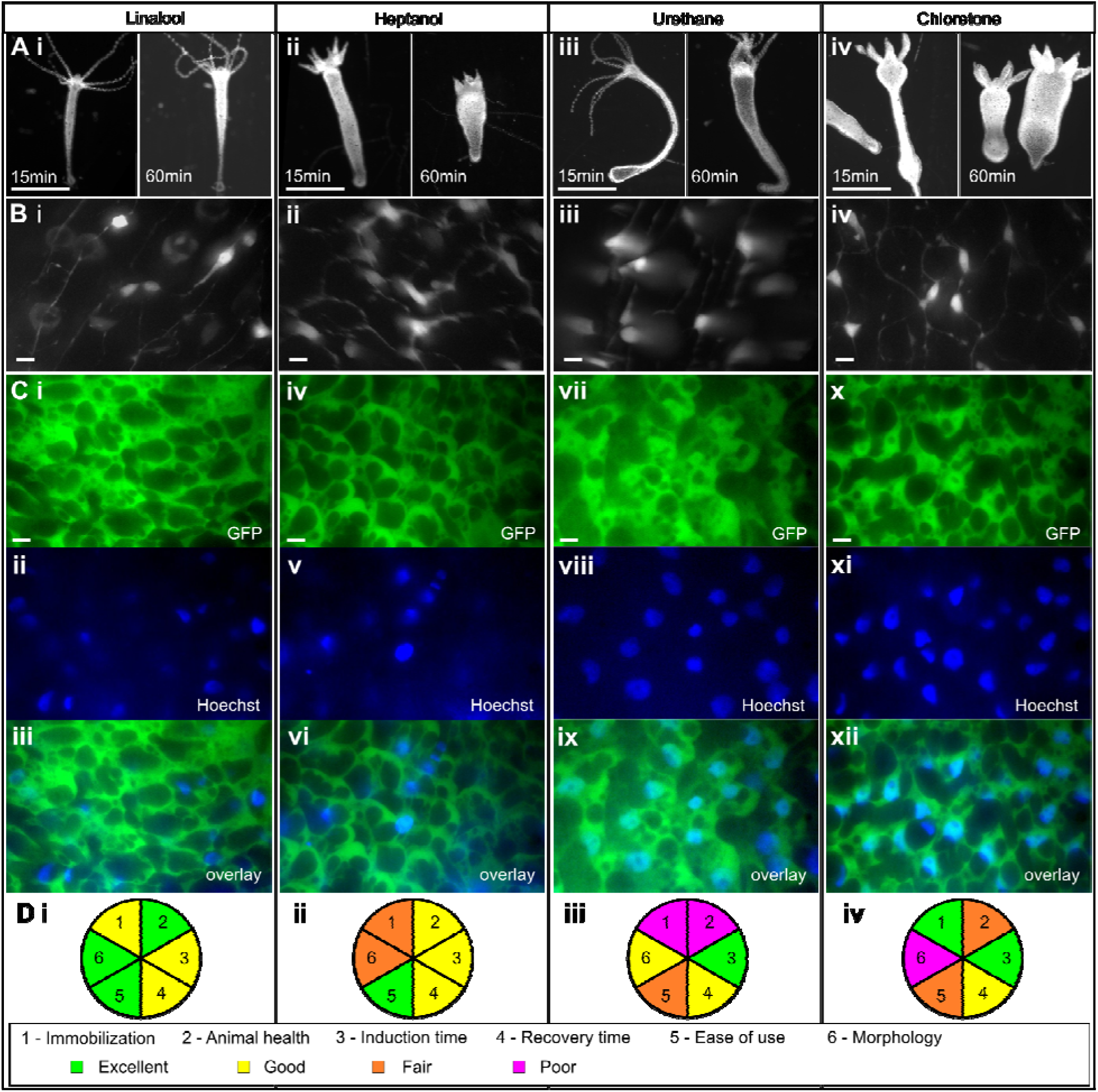
Comparison of various *Hydra* anesthetics. A. Comparisons of the same animal after 15 min and 60 min of anesthetic exposure. i. 1 mM linalool, ii. 0.04% heptanol, iii. 2% urethane, iv. 0.1% chloretone. Scale bar: 1 mm B. Maximum intensity projections of GCaMP6s animals at 60x magnification in each anesthetic. Scale bars: 10 µm C. Maximum intensity projections of two-channel images of watermelon animals stained with Hoechst nuclear dye at 60x magnification. GFP channel, DAPI channel, and merge shown for each anesthetic. Scale bars: 10 µm. D. Overview of the four anesthetics tested, scored on degree of immobilization, animal health following anesthesia, time to induce anesthesia, time to recover from anesthesia, and ease of use.

Because most published studies specified only the concentration of anesthetic used and not the incubation time, we used concentrations that have been reported in the literature to be effective for the different anesthetics and measured induction and recovery times for direct comparison to linalool. Overall, the times were fairly similar. Linalool’s induction time (9min, (6, 9)) was similar to that of heptanol (6min, (4, 9)), but significantly longer than those of urethane (5min, (4, 6)) and chloretone (5min, (3, 7)) (Fig. S4B, E). However, as the average induction time for linalool is below 10 minutes, this is acceptable for routine use. Recovery times were statistically similar between all anesthetics, with most polyps resuming normal activity within 10-20 min post-exposure (Fig. S4C, F).

Finally, we compared the effects of long-term exposure to the different anesthetics. First, we tested a 3 day exposure to anesthetic without changes of medium, as would be necessary for long term immobilization for continuous imaging. 1mM linalool does not negatively affect intact polyps (Fig. 6), but immobilizes them sufficiently to allow continuous imaging of cellular processes in the intact animal over the course of 3 days. In contrast, all polyps disintegrated within 24h upon continuous exposure in 2% urethane (Fig. S5A). Under the same conditions, chloretone caused disintegration in only a small fraction of animals (Fig. S5A). However, the animals that survived the 3-day chloretone treatment without solution exchange had a fairly normal morphology and pinch response, potentially due to a developed tolerance, as previously suggested (Kepner & Hopkins 1938). Heptanol was not lethal to *Hydra* over 3 days (Fig. S5A), but similar to chloretone the animals regained normal morphology and pinch response by the third day.

Subsequently we tested a 3 day incubation with media changes every 24h, to determine whether performance could be improved by constant refreshing of the anesthetic. Urethane was excluded from this experiment due to its rapid lethality. Linalool had no impact on survival. Significant differences were observed in the cases of chloretone and heptanol. Chloretone caused disintegration of all animals by 48h, whereas heptanol killed ∼85% of the treated animals by 72h. (Fig. S5B)

Finally, we also compared the performance of these various anesthetics for single and dual channel high magnification fluorescent live imaging of GCaMP6s (Fig. 7B) and WM animals labeled with Hoechst (Fig. 7C), respectively. While we did not observe a big difference in stability and image quality across the different conditions as observed earlier when comparing to animals in HM (Fig. 3), linalool or chloretone exposed animals allowed us to get slightly better results than animals exposed to heptanol or urethane. For both linalool and chloretone, we were able to obtain crisp images of neuronal processes without blurring (Fig. 7B i, iv) and the nuclear staining was better co-localized with the cells (Fig. 7C i, iv). To summarize our direct comparison of the various anesthetics, linalool has the best overall performance (Fig. 7D), although chloretone provided better images using short-term high-resolution fluorescence microscopy.

## Discussion

Our results show that linalool is a fast-acting, reversible anesthetic for *Hydra.* It is non-toxic and simple to use, and its pleasant smell makes working with it an enjoyable experience. Incubation in 1 mM linalool does not completely immobilize the animal, as we have observed mouth opening and feeding (Fig. 1F, 4A). The ability of polyps to open their mouths and feed in the absence of a pinch response suggests that linalool affects primarily the body column. As both spontaneous and mechanically induced contractions were absent in linalool-exposed polyps, linalool must affect epitheliomuscular cells.

In terms of applications, a 10 min incubation in 1 mM linalool significantly decreased polyp movement, allowing for fine surgical manipulations with superior precision, efficiency, and long-term success compared to their execution in HM (Fig. 2). Additionally, we achieved improved fluorescent imaging when compared to HM and were able to acquire good quality single- and multi-channel fluorescent z-stacks and time lapse movies (Fig.4 and Supplemental Movies). Furthermore, we showed that linalool enables repeated short-term imaging of the same specimen over the course of days, allowing us to visualize the dynamics of graft development and head regeneration in individuals (Fig. 2D and Fig. 5A). We were able to achieve fluorescent imaging with sub-cellular resolution (Fig. 3), which suggests that one could study cellular migration processes over the course of days.

When compared to other currently used anesthetics, 1 mM linalool is superior in terms of ease of preparation, handling, and disposal. Linalool and heptanol are both alcohols and supplied as liquids. Working concentrations are easily made up fresh within minutes by pipetting the adequate amount of stock solution. Because heptanol has a strong smell, however, preparation in the fume hood may be preferred. Urethane and chloretone are powders and therefore the preparation of stock solutions requires more time and safety precautions, such as working in a fume hood. In terms of overall toxicity, linalool is the least harmful substance and considered non-toxic at the concentrations employed here (Letizia et al. 2003). Chloretone is also comparably non-toxic at the concentrations used here (Nordt 1996), whereas urethane is a known carcinogen (Tuveson & Jacks 1999; Salaman & Roe 1953; Sakano et al. 2002). Heptanol is considered to have aquatic toxicity (Slooff et al. 1983), has been shown to be a teratogen (Bernardini et al. 1994), and causes abnormal patterning phenotypes, such as two-headed animals in freshwater planarians upon long-term low-dose exposure (Nogi & Levin 2005). Thus, linalool provides a clear advantage in terms of ease of use and lack of toxicity.

Linalool also has an advantage in anesthetized animal morphology (Fig 7A). Because animals extend in linalool similar to what is observed in urethane, precision cuts and grafting experiments are facilitated (Fig. 2); in contrast, animals immobilized in heptanol or chloretone appear contracted and misshapen (Fig.7A ii, iv). As grafting requires precise cuts and manipulations that are most easily executed on an evenly extended animal, chloretone and heptanol are suboptimal for such applications. 1 mM linalool outperforms 2% urethane and 0.04% heptanol for short-term high-magnification fluorescence imaging applications, but produces a slightly inferior image quality when compared to 0.1% chloretone (Fig. 6 B,C). However, the demonstrated lack of cell damage or other harm to the animal may represent an advantage for repeated imaging or for particularly sensitive experiments.

In terms of long-term applications, we find that a 3-day continuous exposure without media exchange is lethal in urethane within 24 h, partially lethal in chloretone, and harmless in linalool and heptanol. Surviving chloretone and heptanol-treated animals showed normal morphology and a pinch response. The observed detrimental effect on animal health of urethane may be due to an overly broad mechanism of action that impacts other aspects of the animal’s biology Urethane has been shown to act in *Hydra* by reversing the sodium polarity across the cell membrane and to lead to structural damage (Macklin 1976). While chloretone has been proposed to act directly on nerves (Kepner & Hopkins 1938), its mechanism of action in *Hydra* remains unclear. It is possible that the gross anatomical changes that are observed in exposure to chloretone cause functional problems that ultimately cause death. Heptanol is a gap junction blocker that effectively blocks ectodermal epithelial cell-cell communication in the body column at 0.04% v/v (Takaku et al. 2015). As a small alcohol, its effect may be lost over long incubations due to its volatility. Exchanging the media every 24h drastically changed the outcome of incubation in chloretone and heptanol, with all chloretone-treated and the majority of heptanol-treated animals dying within 3 days. This result suggests that the survival and loss of anesthesia seen in animals incubated 72h in heptanol or chloretone without medium changes is due to evaporation or degradation of the chemical, and that continuous exposure to active concentrations is toxic to the animals. In summary, these data suggest that urethane, chloretone, and heptanol cannot be used for continuous 3-day exposure and long-term imaging. Thus, linalool is the only viable option among the four anesthetics tested for long-term experiments.

In contrast to intact polyps, regenerating body columns that were continuously exposed to 1 mM linalool over 3 days showed negative health effects, including abnormal morphology (Fig. 6E ii), suppressed regeneration (Fig. 6D), and loss of cells. Both head and foot regeneration were delayed (Fig. 6D and Fig. S2). Affected animals healed their wounds, but did not develop the structures associated with the missing body part – decapitated animals did not form tentacles or hypostomes, and animals lacking feet did not regain a peduncle or the ability to adhere to the substrate. Regeneration could be rescued by transferring the regenerating animals back to HM after 3 days of linalool exposure (Fig. 6E). While these findings prevent the use of linalool for continuous long-term imaging of regeneration, they indicate that linalool could potentially be a useful tool for regeneration studies if the mechanism of action can be elucidated.

Because nerve-free animals in linalool do not show delayed regeneration (Fig. S3), these data suggest that nerve or interstitial cells are the target for the regeneration effect. The precise role of the interstitial cell lineage in regeneration and morphogenesis is unknown. It has been shown that nerve-free *Hydra* are fully capable of regeneration and budding (Marcum & Campbell 1978). Marcum and Campbell propose several possible explanations for this observation – 1. that nerve cells are not involved in development, 2. that nerve cells modulate developmental processes initiated by epithelial cells, 3. that nerve cells play an essential role in patterning but that their absence can be compensated for, or 4. that nerve and epithelial cells both have critical but overlapping roles in development. Head regeneration is delayed in *Hydra* treated with double-stranded RNA from a gene encoding a neuronal progenitor marker, leading to the idea that neurons are required for head regeneration (Miljkovic-Licina et al. 2007). The authors take this result to support the third possibility laid out by Marcum and Campbell – that neurons are critical for regeneration, but that in their complete absence nerve-free animals can employ an alternate pathway. Our finding that linalool prevents regeneration in wild type animals while having no effect on nerve-free animals supports this idea that neuronal signals play an important role for head regeneration under normal circumstances. It will be exciting to dissect this relationship between nerve signaling and axial patterning. One possible starting point for investigation is linalool’s known mechanism of action in other systems.

Linalool has been found to modify nicotinic receptors at neuromuscular junctions in rodents, leading to modulated acetylcholine release (Re et al. 2000) and to inhibit glutamatergic signaling in the central nervous system (Elisabetsky et al. 1995). While it is unclear whether the mechanism of anesthesia in *Hydra* is the same as that in rodents, the cellular machinery targeted is sufficiently conserved that this is a possibility. Full characterization of neurotransmitters and their receptors in *Hydra* has proven elusive thus far but there is still a broad base of evidence for glutamatergic and cholinergic signaling in *Hydra*. *Hydra* has been shown to possess GABA receptors (Pierobon et al. 1995), and to have specific glutamate-binding abilities likely corresponding to at least two types of glutamate receptors (Bellis et al. 1991). GABA, glutamate, and their agonists and antagonists have been shown to influence behaviors such as contraction bursts (Kass-Simon et al. 2003), as well as nematocyst activity (Kass-Simon & Scappaticci 2004). Similarly, *Hydra* homogenate was found to contain an enzyme that hydrolyzes acetylcholine (Eržen & Brzin 1978). Nicotinic acetylcholinesterase antagonists were found to decrease contraction bursts while a muscarinic acetylcholinergic antagonist increased them (Kass-Simon & Passano 1978). A cDNA sequence for acetylcholinesterase has also been cloned, though its expression and localization have not been confirmed (Takahashi & Hamaue 2010). Thus, linalool’s mechanism of action may be conserved between *Hydra* and rodents, though further mechanistic studies will be needed to test this hypothesis.

In summary, linalool offers a range of advantages over other available anesthetics by enabling new applications such as long term or repeated imaging while also being usable as a pre-fixation relaxant in the same way as current options. Linalool’s lack of toxicity to both *Hydra* and researchers and the ease of use and preparation compared to current anesthetics render it an attractive tool for *Hydra* experimentation in the teaching setting. In particular, linalool makes grafting experiments that can provide fundamental insights into regeneration and biological patterning accessible to students with no previous experience with *Hydra*.

## Conclusion

The advent of modern biological tools to generate and manipulate transgenic *Hydra* lines has sparked a new interest in this fascinating model system because it allows for unprecedented *in vivo* studies to dissect the mechanisms underlying regeneration and animal behavior. Here we introduce linalool as a powerful anesthetic to reliably and reversibly relax *Hydra* tissue or whole animals and demonstrate its usefulness for *in vivo* tissue manipulation and short-term high-resolution fluorescent imaging. Linalool outperforms any other currently used anesthetic in ease of use, lack of toxicity to both the animal and the researcher, and overall performance as a reversible anesthetic.

## Acknowledgements

The authors thank Trevor Rowe for the creation of the “Frank” animals, Connor Keane for help with the experiments and data analysis, Dr. Robert Steele and Dr. Danielle Ireland for discussion and comments on the manuscript, Dr. Nick Kaplinsky for the Syto 60 stain, and Dr. Rafael Yuste, Dr. Christophe Dupre, and Dr. Alison Hanson for the GCaMP6s and A10 strains.

## References

Ando, H. et al., 1989. Pattern formation in hydra tissue without developmental gradients. Developmental Biology, 133(2), pp.405–414.

Aprotosoaie, A.C. et al., 2014. Linalool: A review on a key odorant molecule with valuable biological properties. Flavour and Fragrance Journal, 29(4), pp.193–219.

Badhiwala, K.N. et al., 2018. Microfluidics for electrophysiology, imaging, and behavioral analysis of Hydra †., 18, p.2523.

Bellis, S.L. et al., 1991. Chemoreception in Hydra vulgaris (attenuata): initial characterization of two distinct binding sites for l-glutamic acid. Biochimica et Biophysica Acta (BBA) – Biomembranes, 1061(1), pp.89–94.

Benos, D.J. et al., 1977. Hyposmotic fluid formation in Hydra. Tissue and Cell, 9(1), pp.11–22.

Bernardini, G. et al., 1994. Lethality, teratogenicity and growth inhibition of heptanol in Xenopus assayed by a modified frog embryo teratogenesis assay-Xenopus (FETAX) procedure. Science of The Total Environment, 151(1), pp.1–8.

Bode, H. et al., 1973. Quantitative analysis of cell types during growth and morphogenesis in Hydra. Wilhelm Roux Archiv für Entwicklungsmechanik der Organismen, 171(4), pp.269–285.

Bode, H.R., 2012. The head organizer in Hydra. The International Journal of Developmental Biology, 56(6-7–8), pp.473–478.

Bode, H.R., 1996. The interstitial cell lineage of hydra: a stem cell system that arose early in evolution. Journal of cell science, 109, pp.1155–1164.

Boothe, T. et al., 2017. A tunable refractive index matching medium for live imaging cells, tissues and model organisms. eLife – Tools and Resources.

Bosch, T.C.G., 2009. Hydra and the evolution of stem cells. BioEssays, 31(4), pp.478–486.

Bosch, T.C.G., 2007. Why polyps regenerate and we don’t: Towards a cellular and molecular framework for Hydra regeneration. Developmental Biology, 303(2), pp.421–433.

Broun, M. & Bode, H.R., 2002. Characterization of the head organizer in hydra. Development, 129(4).

Browne, E.N., 1909. The production of new hydranths in Hydra by the insertion of small grafts. Journal of Experimental Zoology, 7(1), pp.1–23.

Burnett, A.L. & Diehl, N.A., 1964. The nervous system of hydra. I. Types, distribution and origin of nerve elements. Journal of Experimental Zoology, 157(2), pp.217–226.

Buzgariu, W. et al., 2018. Impact of cycling cells and cell cycle regulation on Hydra regeneration. Developmental Biology, 433(2), pp.240–253.

Campbell, R.D., 1974. Cell Movements in Hydra. American Zoologist, 14(2), pp.523–535.

Campbell, R.D. & David, C.N., 1974. Cell Cycle Kinetics and Development of Hydra attenuata. II. Interstitial cells. Journal of Cell Science, 16(2), pp.349–358.

Carter, J.A. et al., 2016. Dynamics of Mouth Opening in Hydra. Biophysical Journal, 110(5), pp.1191–1201.

Chapman, J.A. et al., 2010. The dynamic genome of Hydra. Nature, 464(7288), pp.592–596.

Cikala, M. et al., 1999. Identification of caspases and apoptosis in the simple metazoan Hydra. Current Biology, 9(17), pp.959–S2.

Cochet-Escartin, O. et al., 2017. Physical Mechanisms Driving Cell Sorting in Hydra. Biophysical Journal, 113(12), pp.2827–2841.

David, C.N., 1973. A quantitative method for maceration of hydra tissue. Wilhelm Roux Archiv für Entwicklungsmechanik der Organismen, 171(4), pp.259–268.

David, C.N. & Campbell, R.D., 1972. Cell cycle kinetics and development of Hydra attenuata. I. Epithelial cells. Journal of Cell Science, 11(2), pp.557–68.

David, C.N. & Murphy, S., 1977. Characterization of interstitial stem cells in hydra by cloning. Developmental Biology, 58(2), pp.372–383.

Dupre, C. & Yuste, R., 2017. Non-overlapping Neural Networks in Hydra vulgaris. Current Biology, 27(8), pp.1085–1097.

Elisabetsky, E., Marschner, J. & Onofre Souza, D., 1995. Effects of linalool on glutamatergic system in the rat cerebral cortex. Neurochemical Research, 20(4), pp.461–465.

Engel, U. et al., 2002. Nowa, a novel protein with minicollagen Cys-rich domains, is involved in nematocyst formation in Hydra. Journal of cell science, 115(Pt 20), pp.3923–34.

Eržen, I. & Brzin, M., 1978. Cholinergic mechanisms in hydra. Comparative Biochemistry and Physiology Part C: Comparative Pharmacology, 59(1), pp.39–43.

Fujisawa, T., 2003. Hydra regeneration and epitheliopeptides. Developmental Dynamics, 226(2), pp.182–189.

Galliot, B. et al., 2018. Non-developmental dimensions of adult regeneration in Hydra. The International journal of developmental biology, 62(6-7–8), pp.373–381.

Gierer, A. et al., 1972. Regeneration of Hydra from Reaggregated Cells. Nature New Biology, 239(91), pp.98–101.

Gierer, A. & Meinhardt, H., 1972. A Theory of Biological Pattern Formation. Kybernetik, 12, pp.30–39.

Glauber, K.M. et al., 2013. A small molecule screen identifies a novel compound that induces a homeotic transformation in Hydra. Development (Cambridge, England), 140(23), pp.4788–96.

Glauber, K.M. et al., 2015. A small molecule screen identifies a novel compound that induces a homeotic transformation in Hydra. Development, 142(11), pp.2081–2081.

Hagstrom, D. et al., 2015. Freshwater Planarians as an Alternative Animal Model for Neurotoxicology. Toxicological Sciences, 147(1), pp.270–285.

Han, S. et al., 2018. Comprehensive machine learning analysis of Hydra behavior reveals a stable basal behavioral repertoire. eLife, 7.

Hausman, R.E. & Burnett, A.L., 1971. The mesoglea of Hydra. IV. A qualitative radioautographic study of the protein component. Journal of Experimental Zoology, 177(4), pp.435–446.

Heldwein, C.G. et al., 2014. S-(+)-Linalool from Lippia alba: Sedative and anesthetic for silver catfish (Rhamdia quelen). Veterinary Anaesthesia and Analgesia, 41(6), pp.621–629.

Hobmayer, B. et al., 2000. WNT signalling molecules act in axis formation in the diploblastic metazoan Hydra. Nature, 407(6801), pp.186–189.

Höferl, M., Krist, S. & Buchbauer, G., 2006. Chirality Influences the Effects of Linalool on Physiological Parameters of Stress. Planta Medica, 72(13), pp.1188–1192.

Hufnagel, L.A., Kass-Simon, G. & Lyon, M.K., 1985. Functional organization of battery cell complexes in tentacles ofHydra attenuata. Journal of Morphology, 184(3), pp.323–341.

Hufnagel, L.A. & Myhal, M.L., 1977. Observations on a Spirochaete Symbiotic in Hydra. Transactions of the American Microscopical Society, 96(3), p.406.

Juliano, C.E., Lin, H. & Steele, R.E., 2014. Generation of Transgenic Hydra by Embryo Microinjection. Journal of Visualized Experiments, 91, p.e51888.

Kass-Simon, G., Pannaccione, A. & Pierobon, P., 2003. GABA and glutamate receptors are involved in modulating pacemaker activity in hydra. Comparative Biochemistry and Physiology – A Molecular and Integrative Physiology, 136(2), pp.329–342.

Kass-Simon, G. & Passano, L.M., 1978. A neuropharmacological analysis of the pacemakers and conducting tissues of Hydra attenuata. Journal of Comparative Physiology? A, 128(1), pp.71–79.

Kass-Simon, G. & Scappaticci, A.A., 2004. Glutamatergic and GABAnergic control in the tentacle effector systems of Hydra vulgaris. Hydrobiologia, 530–531(1–3), pp.67–71.

Kepner, W.A. & Hopkins, D.L.L., 1938. Reactions of Hydra to Chloretone. Journal of Experimental Zoology, 38(c), pp.951–959.

Koizumi, O., 2002. Developmental neurobiology of hydra, a model animal of cnidarians. Canadian Journal of Zoology, 80(10), pp.1678–1689.

Lenhoff, H.M. & Brown, R.D., 1970. Mass culture of hydra: an improved method and its application to other aquatic invertebrates. Laboratory Animals, 4(1), pp.139–154.

Lenhoff, S.G. & Lenhoff, H.M., 1986. Hydra and the Birth of Experimental Biology – 1744,

Letizia, C.. et al., 2003. Fragrance material review on linalool. Food and Chemical Toxicology, 41(7), pp.943–964.

Linck, V. de M. et al., 2009. Inhaled linalool-induced sedation in mice. Phytomedicine, 16(4), pp.303–307.

Livshits, A. et al., 2017. Structural Inheritance of the Actin Cytoskeletal Organization Determines the Body Axis in Regenerating Hydra. Cell Reports, 18(6), pp.1410–1421.

Lommel, M. et al., 2017. Genetic knockdown and knockout approaches in Hydra. bioRxiv.

Loomis, W.F., 1955. Glutathione control of the specific feeding reactions of hydra. Annals of the New York Academy of Sciences, 62(9 Gluathione Co), pp.211–227.

Macklin, M., 1976. The effect of urethan on hydra. The Biological bulletin, 150(3), pp.442–52.

MacWilliams, H.K., 1983. Hydra transplantation phenomena and the mechanism of Hydra head regeneration: II. Properties of the head activation. Developmental Biology, 96(1), pp.239–257.

Marcum, B.A. & Campbell, R.D., 1978. Development of Hydra lacking nerve and interstitial cells. Journal of Cell Science, 29(1).

Martin, V.J. et al., 1997. Embryogenesis in hydra. The Biological bulletin, 192(3), pp.345–63.

Miljkovic-Licina, M. et al., 2007. Head regeneration in wild-type hydra requires de novo neurogenesis. Development (Cambridge, England), 134(6), pp.1191–201.

Münder, S. et al., 2013. Notch-signalling is required for head regeneration and tentacle patterning in Hydra. Developmental Biology, 383(1), pp.146–157.

Nogi, T. & Levin, M., 2005. Characterization of innexin gene expression and functional roles of gap-junctional communication in planarian regeneration. Developmental Biology, 287(2), pp.314–335.

Nordt, S.P., 1996. Chlorobutanol toxicity. The Annals of pharmacotherapy, 30(10), pp.1179–80.

Noro, Y. et al., 2019. Regionalized nervous system in Hydra and the mechanism of its development. Gene Expression Patterns, 31, pp.42–59.

Otto, J.J. & Campbell, R.D., 1977. Budding inHydra attenuata: Bud stages and fate map. Journal of Experimental Zoology, 200(3), pp.417–428.

Petersen, H.O. et al., 2015. A Comprehensive Transcriptomic and Proteomic Analysis of Hydra Head Regeneration. Molecular Biology and Evolution, 32(8), pp.1928–1947.

Pierobon, P. et al., 1995. Biochemical and functional identification of GABA receptors in Hydra vulgaris. Life Sciences, 56(18), pp.1485–1497.

Rand, H.W., Bovard, J.F. & Minnich, D.E., 1926. Localization of Formative Agencies in Hydra. Proceedings of the National Academy of Sciences of the United States of America, 12(9), pp.565–70.

Re, L. et al., 2000. Linalool modifies the nicotinic receptor-ion channel kinetics at the mouse neuromuscular junction. Pharmacological Research, 42(2), pp.177–181.

Rentzsch, F., Hobmayer, B. & Holstein, T.W., 2005. Glycogen synthase kinase 3 has a proapoptotic function in Hydra gametogenesis. Developmental Biology, 278(1), pp.1–12.

Rodenak-Kladniew, B. et al., 2018. Linalool induces cell cycle arrest and apoptosis in HepG2 cells through oxidative stress generation and modulation of Ras/MAPK and Akt/mTOR pathways. Life Sciences, 199(February), pp.48–59.

Sakano, K. et al., 2002. Metabolism of carcinogenic urethane to nitric oxide is involved in oxidative DNA damage. Free Radical Biology and Medicine, 33(5), pp.703–714.

Salaman, M.H. & Roe, F.J., 1953. Incomplete carcinogens: ethyl carbamate (urethane) as an initiator of skin tumour formation in the mouse. British journal of cancer, 7(4), pp.472–81.

Schindelin, J. et al., 2012. Fiji: an open-source platform for biological-image analysis. Nature Methods, 9(7), pp.676–682.

Shimizu, H. et al., 2002. Epithelial morphogenesis in hydra requires de novo expression of extracellular matrix components and matrix metalloproteinases. Development, 129(6), p.1521 LP-1532.

Shimizu, H., Koizumi, O. & Fujisawa, T., 2004a. Three digestive movements in Hydra regulated by the diffuse nerve net in the body column. Journal of Comparative Physiology A, 190(8), pp.623–630.

Shimizu, H., Koizumi, O. & Fujisawa, T., 2004b. Three digestive movements in Hydra regulated by the diffuse nerve net in the body column. Journal of Comparative Physiology A, 190(8), pp.623–630.

Shimizu, H. & Sawada, Y., 1987. Transplantation phenomena in hydra: Cooperation of position-dependent and structure-dependent factors determines the transplantation result. Developmental Biology, 122(1), pp.113–119.

Shimizu, H., Sawada, Y. & Sugiyama, T., 1993. Minimum Tissue Size Required for Hydra Regeneration. Developmental Biology, 155(2), pp.287–296.

Siebert, S. et al., 2018. Stem cell differentiation trajectories in Hydra resolved at single-cell resolution. bioRxiv, p.460154.

Slooff, W., Canton, J.H. & Hermens, J.L.M., 1983. Comparison of the susceptibility of 22 freshwater species to 15 chemical compounds. I. (Sub)acute toxicity tests. Aquatic Toxicology, 4(2), pp.113–128.

Smith, K.M., Gee, L. & Bode, H.R., 2000. HyAlx, an aristaless-related gene, is involved in tentacle formation in hydra. Development, 127(22).

Steele, R.E., 2002. Developmental Signaling in Hydra: What Does It Take to Build a “Simple” Animal? Developmental Biology, 248(2), pp.199–219.

Sugiyama, T. & Fujisawa, T., 1978. Genetic analysis of developmental mechanisms in Hydra. II. Isolation and characterization of an interstitial cell-deficient strain. Journal of cell science, 29, pp.35–52.

Takahashi, T. & Hamaue, N., 2010. Molecular characterization of Hydra acetylcholinesterase and its catalytic activity. FEBS Letters, 584(3), pp.511–516.

Takaku, Y. et al., 2015. Innexin gap junctions in nerve cells coordinate spontaneous contractile behavior in Hydra polyps. Scientific Reports, 4(1), p.3573.

Technau, U. et al., 2003. Arrested apoptosis of nurse cells during Hydra oogenesis and embryogenesis. Developmental Biology, 260(1), pp.191–206.

Technau, U. et al., 2000. Parameters of self-organization in Hydra aggregates. Proceedings of the National Academy of Sciences of the United States of America, 97(22), pp.12127–31.

Thévenaz, P., Ruttimann, U.E. & Unser, M., 1998. A Pyramid Approach to Subpixel Registration Based on Intensity. IEEE Transactions on Image Processing, 7(1), pp.27–41.

Tran, C.M. et al., 2017. Generation and long-term maintenance of nerve-free Hydra. J. Vis. Exp., 125, p.e56115.

Tuveson, D.A. & Jacks, T., 1999. Modeling human lung cancer in mice: similarities and shortcomings. Oncogene, 18(38), pp.5318–5324.

Wittlieb, J. et al., 2006. Transgenic Hydra allow in vivo tracking of individual stem cells during morphogenesis. Proceedings of the National Academy of Sciences.

Yao, T., 1945. Studies on the Organizer Problem in Pelmatohydra Oligactis. Journal of Experimental Biology, 21(3–4).

