## Supplementary information for "Linalool acts as a fast and reversible anesthetic in *Hydra*"

**Supplementary Material**

**Supplementary Figures**

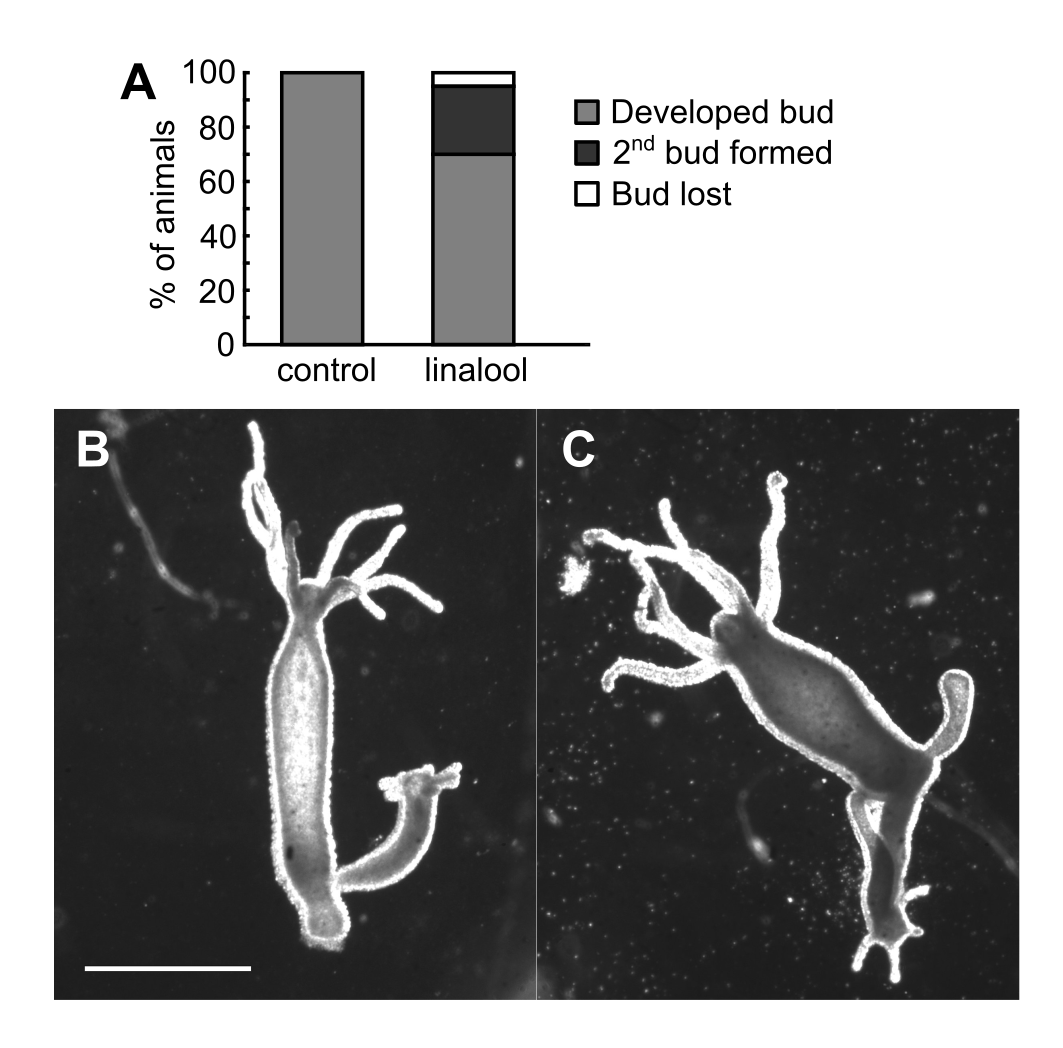

**Fig. S1:** Linalool does not impact budding rate. A. Bud development in budding animals incubated for 3d in HM or 1 mM linalool, n=20 animals per condition. B. Representative image of animal with fully developed bud. C. Representative image of animal with two buds. Scale bar 1 mm.

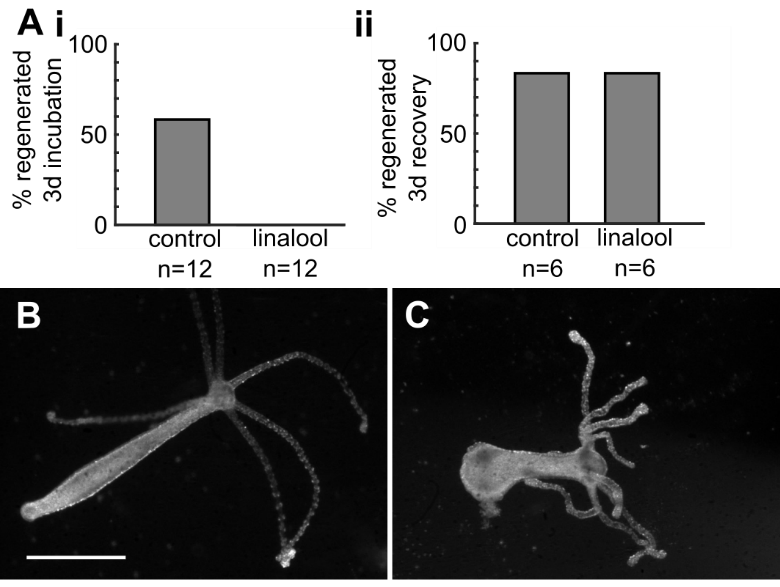

**Fig. S2:** Linalool inhibits foot regeneration. A. i. 3d incubation in linalool prevents foot regeneration, but ii. phenotype is rescued after 3d recovery in HM. B. Polyp incubated 3d in HM after foot amputation. C. Polyp incubated 3d in 1mM linalool after foot amputation. Scale bar 1 mm.

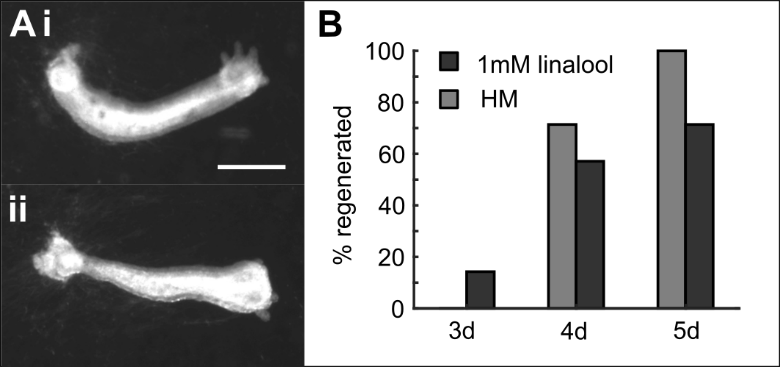

**Fig. S3:** Nerve-free *Hydra* are not negatively impacted by long incubations in linalool. A. Representative images of polyps regenerating their heads in i. HM and ii. linalool after 96 h incubation. Scale bar 1 mm. B. Head regeneration over time.

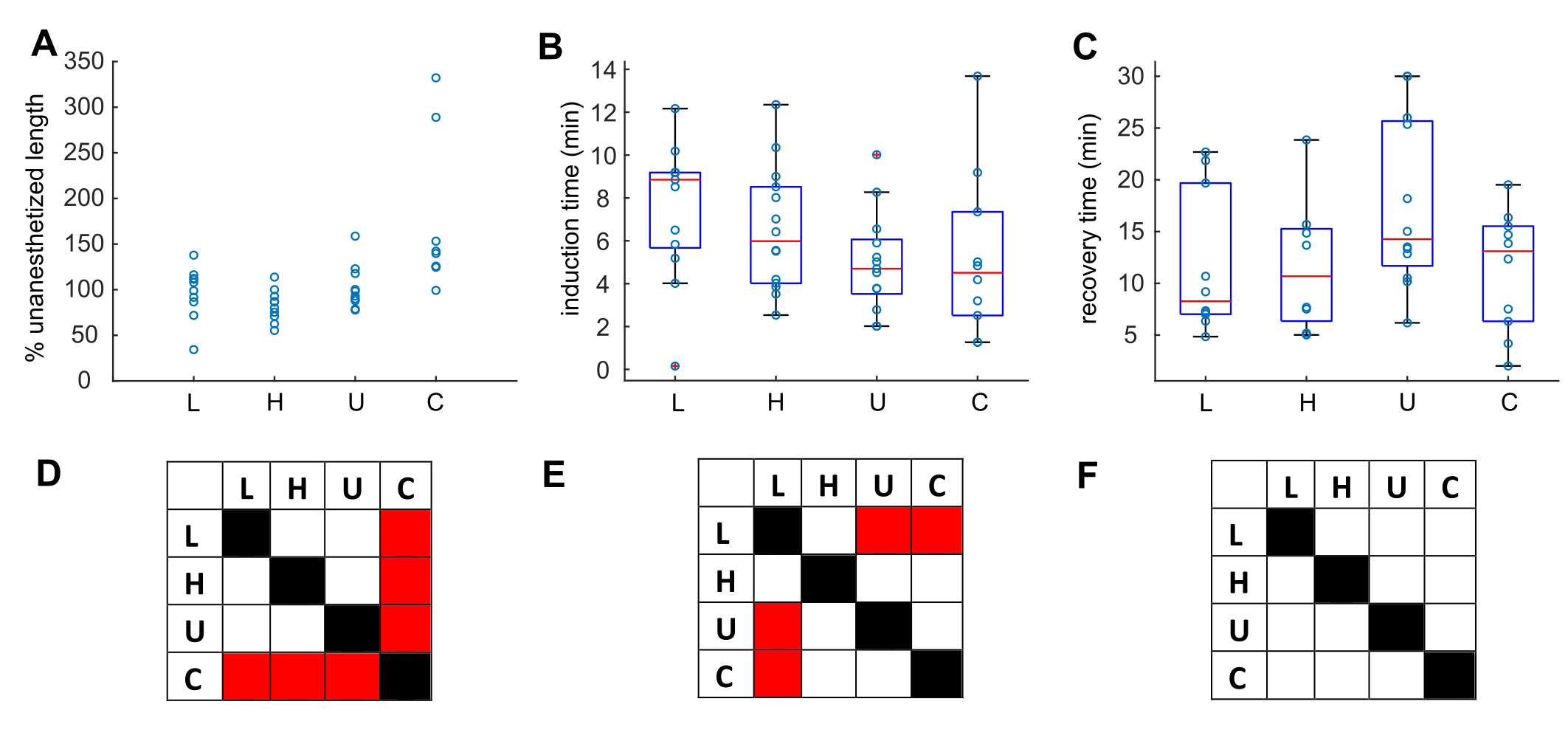

**Fig. S4:** *Hydra* response to 1mM linalool (L), 0.04% heptanol (H), 2% urethane (U), and 0.1% chloretone (C). A. Percent length of anesthetized *Hydra* polyps compared to their natural state. N=10 animals per condition. B. Induction times across 2 technical replicates. Linalool n=13, heptanol n=14, urethane n=13, chloretone n=10. C. Recovery times across 2 technical replicates. Linalool n=10, heptanol n=8, urethane n=12, chloretone n=10. D. Comparison between percent length distributions. Red indicates a statistically significant difference at the p=0.05 level, determined using the Kolmogorov-Smirnov test. E. Comparison between induction time distributions. F. Comparison between recovery time distributions.

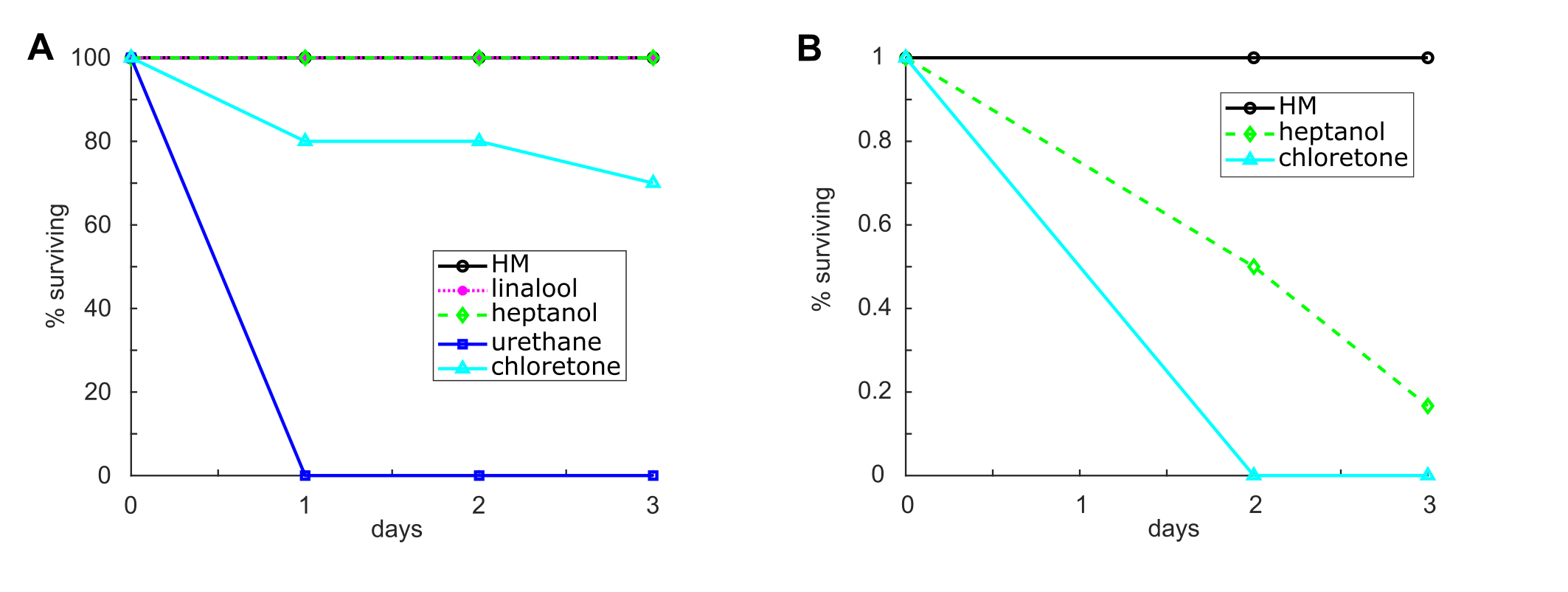

**Fig. S5:** Lethality of three 72h incubation in various anesthetics. A. Incubation without changing media. n=10 animals per condition. Surviving heptanol and chloretone animals had a normal pinch response at 3d. Surviving linalool animals remained anesthetized. B. Incubation with media exchanged every 24h. n=12 animals per condition.

**Supplementary Movies**

**Movie S1.** Time-lapse movie of freely moving GCaMP6s animals in either *Hydra* medium (HM; left) or incubated for at least 10 min in 1mM linalool (LL; right). Experimental details are provided in Methods in the main text and Figure 3A shows a Maximum Intensity Projection of the video. Video playback: 10fps.

**Movie S2.** Time-lapse movie showing a z-stack through the body column of a GCaMP6s animal in either *Hydra* medium (HM; left) or incubated for at least 10 min in 1mM linalool (LL; right). Experimental details are provided in Methods in the main text and Figure 3B shows a single slice and the Maximum Intensity Projection of the video. Video playback: 10fps.

**Movie S3.** Time-lapse movie of a 3-channel acquisition of a watermelon animal stained with Hoechst nuclear dye. The head was mounted as described in Carter *et al.* (1) in 1mM linalool and flushed with 2mM reduced glutathione to trigger a feeding reaction. While the mouth stays closed during recording, one can clearly see the quality of imaging that can be obtained in linalool, allowing for simultaneous imaging of cell shapes and nuclear positions. Video playback: 10fps.
